# Deficiency of gluconeogenic enzyme PCK1 promotes non-alcoholic steatohepatitis progression and fibrosis through PI3K/AKT/PDGF axis activation

**DOI:** 10.1101/2021.01.12.426294

**Authors:** Qian Ye, Yi Liu, Guiji Zhang, Haijun Deng, Xiaojun Wang, Lin Tuo, Chang Chen, Xuanming Pan, Kang Wu, Jiangao Fan, Qin Pan, Kai Wang, Ailong Huang, Ni Tang

## Abstract

Nonalcoholic steatohepatitis (NASH) is a chronic liver disease characterized by hepatic lipid accumulation, inflammation, and progressive fibrosis. We demonstrated that phosphoenolpyruvate carboxykinase 1 (PCK1) plays a central role in NASH progression. Mice with liver *Pck1* deficiency fed a normal diet displayed hepatic lipid disorder and liver injury, whereas fibrosis and inflammation were aggravated in mice fed a NASH diet. Forced expression of PCK1 by adeno-associated virus in the liver ameliorated NASH in mice. PCK1 deficiency stimulated lipogenic gene expression and lipid synthesis. Moreover, loss of hepatic PCK1 activated the RhoA/PI3K/AKT pathway by increasing intracellular GTP levels, increasing secretion of platelet-derived growth factor-AA (PDGF-AA), and promoting hepatic stellate cell activation. Treatment with RhoA and AKT inhibitors or gene silencing of RhoA or AKT1 alleviated NASH progression *in vivo*. Hepatic PCK1 deficiency may be important in hepatic steatosis and fibrosis development through paracrine secretion of PDGF-AA, highlighting a therapeutic strategy for NASH.

## Introduction

Non-alcoholic fatty liver disease (NAFLD) is the most common chronic liver disease worldwide, affecting nearly 25% of US and European adults ^1^. NAFLD is characterized by aberrant lipid accumulation in hepatocytes in the absence of excessive alcohol consumption. Nonalcoholic fatty liver (NAFL) can progress to non-alcoholic steatohepatitis (NASH), a more serious form of liver damage hallmarked by irreversible pathological changes such as inflammation, varying degrees of fibrosis, and hepatocellular damage, which is more likely to develop into cirrhosis and hepatocellular carcinoma ^2^. Although multiple parallel insults, including oxidative damage, endoplasmic reticulum stress, and hepatic stellate cell (HSC) activation, have been proposed to explain the pathogenesis of NASH, the underlying mechanisms remain unclear ^3^.

In gluconeogenesis, glucose is generated from non-carbohydrate substrates, such as glycerol, lactate, pyruvate, and glucogenic amino acids, mainly in the liver, to maintain glucose levels and energy homeostasis. Phosphoenolpyruvate carboxykinase 1 (PCK1) is the first rate-limiting enzyme in gluconeogenesis and converts oxaloacetate to phosphoenolpyruvate in the cytoplasm ^4^. Our previous studies showed that PCK1 deficiency promotes hepatocellular carcinoma progression by enhancing the hexosamine-biosynthesis pathway ^5^ However, PCK1 regulates not only glucose homeostasis but also lipogenesis by activating sterol regulatory element-binding proteins ^6^. Patients lacking PCK1 function present diffuse hepatic macrosteatosis concomitant with hypoglycemia and hyperlactacidemia ^7^. Similarly, mice with reduced *Pck1* expression develop insulin resistance, hypoglycemia, and hepatic steatosis, indicating the important role of PCK1 in regulating both glucose homeostasis and lipid metabolism ^8,9^. However, the precise role of PCK1 in NASH progression is not well-understood.

The phosphoinositide 3-kinase/protein kinase B (PI3K/ATK) pathway plays a critical role in regulating cell growth and metabolism. This pathway is activated in response to insulin, growth factors, energy, and cytokines and, in turn, regulates key metabolic processes such as glucose and lipid metabolism and protein synthesis ^10^. AKT promotes *de novo* lipogenesis primarily by activating sterol regulatory element-binding protein ^11^. PI3K/AKT dysregulation leads to numerous pathological metabolic conditions, including obesity and type 2 diabetes ^12^. NAFLD is characterized by disordered glucose and lipid metabolism in the liver. Although the PI3K/AKT pathway is a key regulator for sensing metabolic stress, its exact role in NAFLD/NASH progression is unclear ^13,14^.

In this study, we explored the role of *Pck1* in a mouse NASH model. We determined the molecular mechanisms underlying disordered lipid metabolism, inflammation, and fibrosis induced by *Pck1* depletion. We also delineated the functional importance of the PI3K/AKT pathway and paracrine secretion of PDGF-AA as its effectors in steatohepatitis, providing a potential therapeutic strategy for treating NASH.

## Results

### PCK1 is downregulated in patients with NASH and mouse models of NASH

To determine whether PCK1 is involved in NAFLD, we first examined hepatic gene expression in a published transcriptome dataset [Gene Expression Omnibus (GEO): GSE126848] containing samples from 14 healthy subjects, 12 patients with obesity, 15 patients with NAFLD, and 16 patients with NASH ^15^. Bioinformatics analysis showed that 32 genes were markedly changed in obesity, NAFLD, and NASH; 12 genes were considerably downregulated and 20 genes were upregulated (Supplementary Fig. 1a–c). Notably, *PCK1* was gradually reduced in patients with obesity, NAFLD, and NASH (Fig. 1a, b). Downregulation of *PCK1* mRNA was also observed in a similar dataset (GSE89632) (Fig. 1b). Moreover, immunohistochemistry (IHC) assays showed that hepatic PCK1 protein levels were significantly lower in patients with NASH than in healthy subjects (Fig. 1c). Similarly, *PCK1* mRNA and protein levels decreased in the liver of mice fed a high-fat diet with drinking water containing fructose and glucose (NASH diet) for 24 weeks (Fig. 1d,e).

**Fig. 1.**
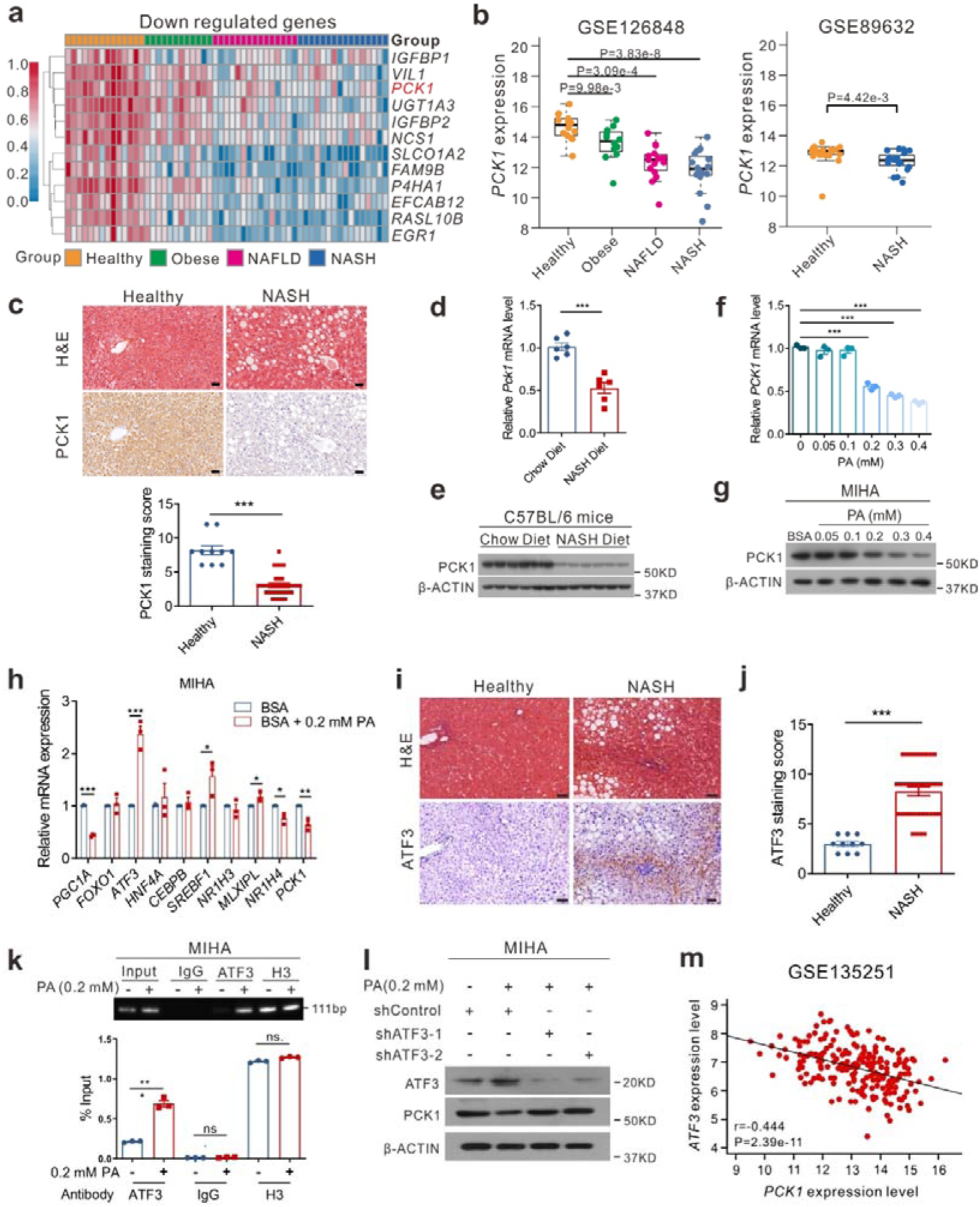
PCK1 is downregulated in patients with NASH and mouse models of NASH. **a** Genes downregulated in patients with obesity (n = 12), NAFLD (n = 15), and NASH (n = 16) from GSE126848 dataset. **b** Relative *PCK1* mRNA levels in GSE126848 and GSE89632 datasets. **c** PCK1 expression in normal individuals and patients with NASH and semi-quantitative analyses of immunohistochemistry (IHC) data (healthy = 10, NASH = 36). Scale bars: 50 µm. **d, e** mRNA (**d**) and protein (**e**) levels of PCK1 in the livers of WT mice fed with chow or NASH diet (n = 6). **F, g** PCK1 mRNA (**f**) and protein (**g**) levels in MIHA cells treated with BSA or PA-BSA. **h** Relative levels of indicated genes in MIHA cells treated with 0.2 mM PA. **i, j** Representative ATF3 expression in normal individuals and patients with NASH (**i**) and semi-quantitative analyses of IHC data (**j**) (healthy = 10, NASH = 36). Scale bars: 50 µm. **k** Chromatin immunoprecipitation assays were performed in MIHA cells with or without PA treatment using an antibody against ATF3, IgG, or H3. **l** Protein levels of PCK1 in MIHA cells infected with either shControl or shATF3 treated with 0.2 mM PA. **m** Correlation analysis of *ATF3* mRNA level with *PCK1* in human NAFLD/NASH liver samples (GSE135251, n = 206). Data are expressed as the mean ± SEM; \**p* < 0.05, \*\**p* < 0.01, \*\*\**p* < 0.001. *p* values obtained via two-tailed unpaired Student’s *t* tests, one-way analysis of variance with Tukey’s post hoc test, or non-parametric Spearman’s test.

Next, palmitic acid (PA) was used to mimic the liver steatosis of patients with NAFLD *in vitro* ^16^. Cell growth was assessed in a CCK8 assay after treatment with different concentrations of PA (Supplementary Fig. 1d). Interestingly, the PCK1 mRNA and protein levels were downregulated in a dose-dependent manner during 24 h PA stimulation (Fig. 1f, g), suggesting that the transcription of PCK1 was inhibited in response to lipid overload. We screened several known regulators of *PCK1* (Supplementary Fig. 1e, f) and found that activating transcription factor 3 (*ATF3*), a transcriptional repressor of *PCK1* ^17^, was upregulated upon PA stimulation (Fig. 1h). Similarly, ATF3 expression was remarkably upregulated in liver samples derived from patients with NASH and NASH model mice (Fig. 1i, j, Supplementary Fig. 1g, h). Chromatin immunoprecipitation assays revealed that the binding of ATF3 to the *PCK1* promoter increased following PA administration (Fig. 1k). ATF3 knockdown restored PCK1 expression in human hepatocytes treated with PA (Fig. 1l). Furthermore, correlation analysis revealed that *ATF3* was negatively correlated with the *PCK1* mRNA level based on the GEO database (GEO: GSE135251) (Fig. 1m). These results indicate that increased lipid intake led to the upregulation of the repressor ATF3, impairing *PCK1* transcription in patients with NASH and mouse models.

### L-KO mice exhibit a distinct hepatic steatosis phenotype

To explore the role of *Pck1* in fatty liver disease, wild-type (WT) and liver-specific *Pck1*-knockout mice (L-KO) mice were fed a chow diet for 24 weeks (Fig. 2a). Hepatic-specific depletion of PCK1 was confirmed by performing immunoblotting (Fig. 2b). Starting at 16 weeks, L-KO mice showed an increased body weight compared with WT mice, however, the results of the glucose tolerance test (GTT) and insulin tolerance test (ITT) did not significantly differ (Fig. 2c). Moreover, significant hepatomegaly and an increased liver weight were observed in L-KO mice (Fig. 2d). Alanine transaminase (ALT) and aspartate transaminase (AST) levels were higher in L-KO mice, indicating liver injury (Fig. 2e). In addition, total triglyceride (TG), total cholesterol (TC), and free fatty acids (FFAs) in the liver tissues and serum were elevated in L-KO mice compared to those in WT mice (Fig. 2f, g). Histochemistry and enzyme-linked immunosorbent assay (ELISA) showed that L-KO mice had prominent hepatic steatosis, increased inflammatory infiltration, and high levels of TNF-α, whereas hepatic fibrosis was not observed (Fig. 2h–j). Additionally, PCK1 deficiency significantly increased the mRNA levels of genes related to fatty acid transport and inflammation, whereas there were no significant changes in fibrosis-related genes in L-KO mice fed the chow diet (Fig. 2k). These data suggest that L-KO mice exhibited a distinct hepatic steatosis phenotype and liver injury even when fed normal chow.

**Fig. 2.**
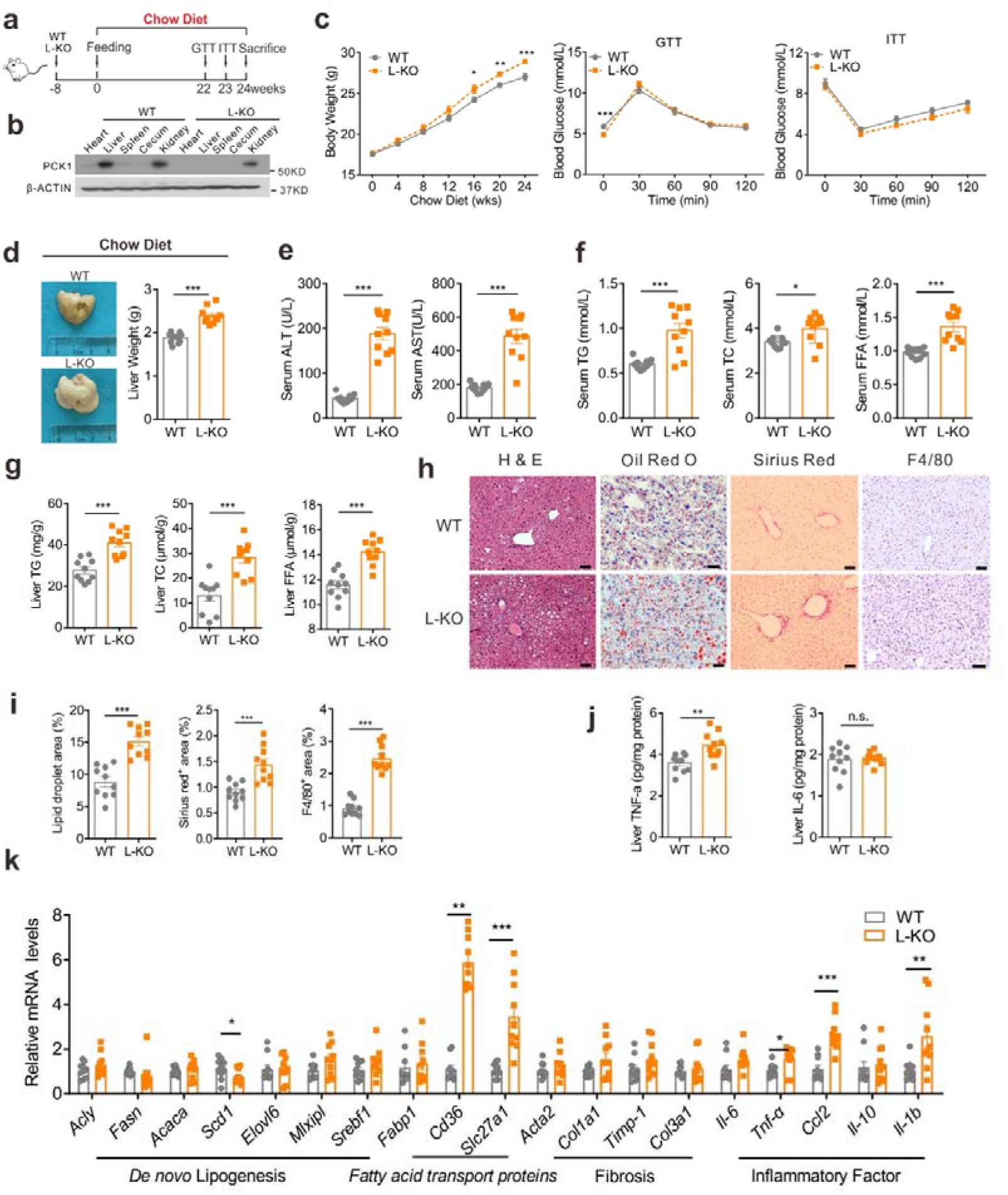
L-KO mice fed chow diet exhibited a distinct hepatic steatosis phenotype. **a** Schematic diagram of the mouse model fed the chow diet, n = 10/group. **b** PCK1 protein expression in WT and L-KO mouse heart, liver, spleen, cecum and kidney were confirmed by immunoblotting. **c** Body weight, glucose tolerance test (GTT), and insulin tolerance test (ITT) were measured in WT and L-KO mice. **d** Representative liver image and liver weight of WT and L-KO mice. **e–g** Determination of alanine aminotransferase (ALT), aspartate aminotransferase (AST), total triglycerides (TG), total cholesterol (TC), and free fatty acid (FFA) levels in the serum or liver tissues. **h** Paraffin-embedded liver sections were stained with hematoxylin and eosin (H&E), Sirius red, and F4/80. Frozen sections stained with Oil Red O. Scale bars: 50 µm. **i** Quantification of liver sections of WT and L-KO mice fed the chow diet. **j** Levels of TNF-α and IL-6 in the liver tissues. **k** Quantitative PCR analysis of liver mRNA expression. Data are expressed as the mean ± SEM; \**p* < 0.05, \*\**p* < 0.01, \*\*\**p* < 0.001; n.s., not significant. *p* values obtained via two-tailed unpaired Student’s *t* tests.

### Hepatic loss of *Pck1* promotes inflammation and fibrogenesis in NASH mice

To explore whether an unhealthy diet could exacerbate pathologic changes in L-KO mice, WT and L-KO mice were fed NASH diet (Fig. 3a) ^18,19^. Starting at 4 weeks, L-KO mice showed significant weight gain (Fig. 3b). The GTT and ITT showed that L-KO mice developed a more severe form of glucose intolerance and insulin resistance compared with those in WT mice (Supplementary Fig. 2a, b). L-KO mice had pale and heavier livers compared with WT mice (Fig. 3c), although there was no significant difference in the liver weight ratio (Supplementary Fig. 2c). Insulin, AST, ALT, TC, TG, and FFAs increased in the serum and liver homogenates of L-KO mice, suggesting more serious liver injury and lipid metabolism disorder (Fig. 3d, Supplementary Fig. 2d, e). Analyses of L-KO liver sections revealed increased fat droplets, more severe fibrosis, and greater macrophage infiltration (Fig. 3e, Supplementary Fig. 2f). Furthermore, L-KO mice exhibited higher NAFLD activity scores (NAS score) and higher TNF-α and IL-6 levels (Fig. 3f, g). In addition, the expression of inflammatory factors, lipogenic enzymes, and fibrogenesis-associated genes was upregulated in L-KO mice (Supplementary Fig. 2g). In summary, mice lacking hepatic *Pck1* showed substantial liver inflammation and fibrosis after being fed the NASH diet.

**Fig. 3.**
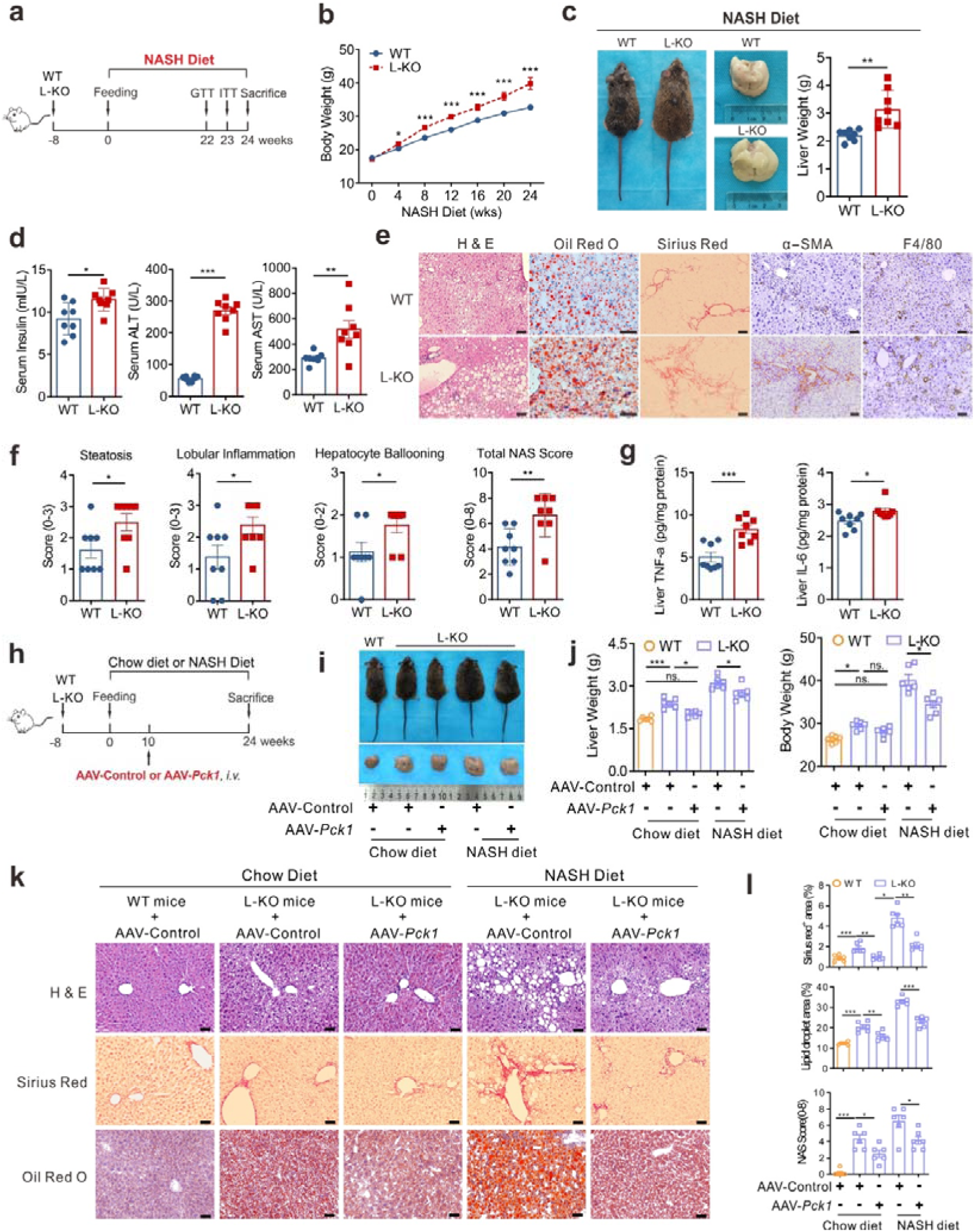
PCK1 ablation accelerates inflammation and fibrogenesis in NASH model. **a** Schematic diagram of mouse model fed the NASH diet. **b** Body weight was measured in WT and L-KO mice. **c** Representative gross liver morphology, whole body photo, and liver weight. **d** Serum levels of insulin, alanine aminotransferase (ALT), and aspartate aminotransferase (AST) were measured. **e** Paraffin-embedded liver sections were stained with hematoxylin and eosin (H&E), Sirius red, α-SMA immunostaining, and F4/80 immunostaining. Frozen sections stained with Oil Red O. Scale bars: 50 µm. **f** Normalized activity scores (NAS) for each group. **g** Levels of TNF-α and IL-6 in the liver tissues detected using enzyme-linked immunosorbent assay (ELISA). **h** Schematic showing the administration protocol for AAV8-TBG-*control* or AAV8-TBG-*Pck1* in WT and L-KO mice for experiments shown in **i**–**l**, n = 6/group. **i** Representative whole-body photo and gross liver morphology. **j** Liver weight and body weight. **k** Paraffin-embedded liver sections were stained with H&E, Sirius red immunostaining and Oil Red O staining. Scale bars: 50 µm. **l** Quantifications of Sirius red staining, Oil Red O staining, and NAS. Data are expressed as the mean ± SEM; \**p* < 0.05, \*\**p* < 0.01, \*\*\**p* < 0.001. *p* values obtained via two-tailed unpaired Student’s *t* tests.

### AAV-mediated restoration of hepatic PCK1 alleviates the NASH phenotype in *Pck1*-deficient mice

We then investigated whether adeno-associated virus (AAV)-based *Pck1* replacement therapy could reverse ongoing liver derangement, which is typically observed in patients with NASH. After 10 weeks of chow or NASH diet feeding, WT and L-KO mice were injected through the tail vein with AAV serotype 8 (AAV8) vector expressing *Pck1* under the control of a liver-specific promoter (thyroxine-binding globulin, TBG), AAV8-TBG-*Pck1* or AAV8-TBG-*control* (Fig. 3h). The expression of PCK1 in various tissues was identified by Western blotting (Supplementary Fig. 2h). Interestingly, mice with PCK1 re-expression showed a lower liver weight, body weight, serum liver enzymes, and serum lipid contents (Fig. 3i, j, Supplementary Fig. 2i, j). Moreover, lipid deposition, inflammation, and fibrosis significantly improved in L-KO mice injected with AAV8-TBG-*Pck1* (Fig. 3k, l). Hepatic gene expression analyses indicated that the expression of genes involved in inflammation and liver fibrosis was greatly attenuated by PCK1 restoration in L-KO mice (Supplementary Fig. 2k). Overall, these data support that forced PCK1 expression in the liver protects against NASH in mice.

### Transcriptomic and metabolomics analyses confirmed that the loss of *Pck1* promotes hepatic lipid accumulation

To comprehensively investigate the role of *Pck1* deficiency in NASH development, we performed RNA-seq analysis of liver samples from L-KO and WT mice fed the normal chow or NASH diet for 24 weeks. Gene Ontology analysis indicated that lipid metabolic processes were remarkably upregulated in L-KO mice fed the NASH diet (Fig. 4a). The volcano plot showed that genes involved in fatty acid uptake, such as solute carrier family 27 member 1 (*Slc27a1*) and fatty acid translocase (*Cd36*), and lipid droplet synthesis, such as cell death-inducing DFFA like effector C (*Cidec*) and cell death-inducing DFFA like effector A (*Cidea*), were upregulated in response to the NASH diet (Fig. 4b). Gene Set Enrichment Analysis (GSEA) revealed that the PPAR signaling pathway was prominently upregulated in L-KO mice fed either diet (Fig. 4c, Supplementary Fig. 3a, b). Several genes selected from the dataset were independently validated by quantitative polymerase chain reaction (qPCR) and immunoblotting and found to be significantly overexpressed in L-KO mice (Fig. 4d, e). Furthermore, genes involved in the glycerol 3-phosphate (G3P) pathway were upregulated in L-KO mice (Fig. 4f). Metabolomics analysis showed that compared with WT mice fed the NASH diet, L-KO mice had significantly higher G3P and PA levels (Fig. 4g, h). As G3P is a substrate for TG synthesis and PA is a key intermediate metabolite in *de novo* lipogenesis, *Pck1* ablation may promote substrate accumulation for lipid synthesis.

**Fig. 4.**
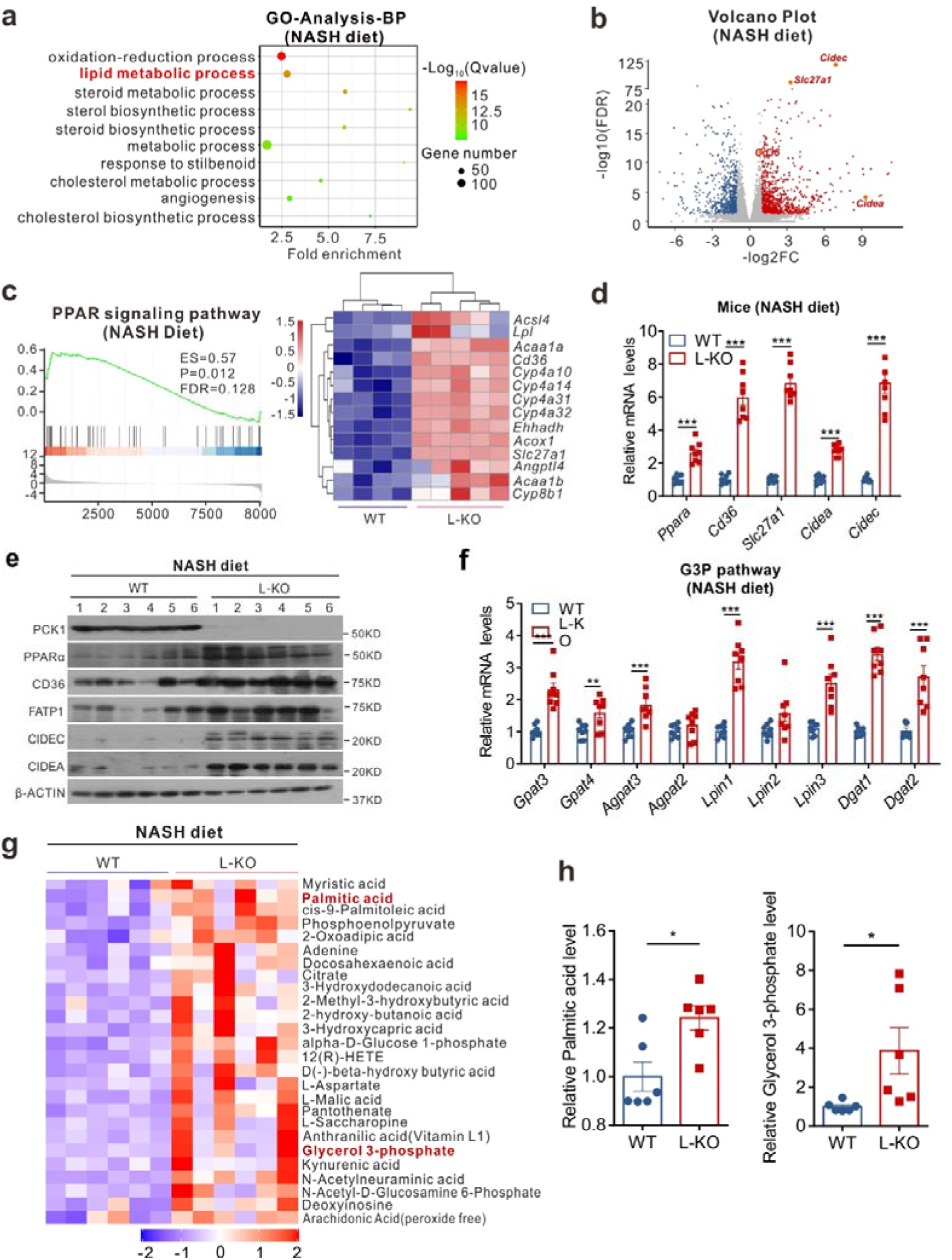
Loss of PCK1 promotes lipid accumulation according to transcriptome and metabolome analyses. RNA sequencing was performed on the livers of WT (n = 4) and L-KO (n = 5) mice fed the NASH diet. **a** Gene Ontology analysis of all significantly changed genes in top 10 biological processes. **b** Volcano plot representation of significantly up- and downregulated genes. **c** Gene Set Enrichment Analysis plot (left) of enrichment in “PPAR signaling pathway” signature; heatmap (right) of significantly upregulated PPAR target genes. **d, e** Quantitative PCR (**d**) and immunoblot (**e**) analysis of indicated genes or protein expression in mouse liver tissues. **f** Relative mRNA expression of key genes in G3P pathway (n = 8). **g** Upregulated metabolites detected by untargeted metabolomics (n = 6). **h** Relative level of G3P and PA in mouse liver tissues (n = 6). Data are expressed as the mean ± SEM; \**p* < 0.05, \*\**p* < 0.01, \*\*\**p* < 0.001. *p* values obtained via two-tailed unpaired Student’s *t* tests.

To further examine the function of PCK1 in steatosis *in vitro*, we overexpressed (*PCK1*-OE) using the AdEasy adenoviral vector system and knocked out PCK1 (*PCK1*-KO) using the CRISPR-Cas9 system in human hepatocytes (Supplementary Fig. 3c, d). We found that *PCK1*-OE attenuated the accumulation of lipid droplets, whereas *PCK1*-KO facilitated lipid accumulation (Supplementary Fig. 3e, f). Collectively, these results suggest that hepatic *Pck1* deficiency leads to lipid accumulation by promoting the expression of lipogenic genes and accumulation of substrates related to lipid synthesis (Supplementary Fig. 3g).

### Hepatic *Pck1* deficiency leads to HSC activation via PI3K/AKT pathway

RNA-seq analysis indicated that the PI3K/AKT pathway was specifically activated in L-KO mice fed the NASH diet (Fig. 5a, b). Immunoblotting revealed p-AKT (S473) and p-AKT (T308), which are two activated forms of AKT, and downstream c-MYC were significantly upregulated in both the livers and primary hepatocytes of L-KO mice fed the NASH diet (Fig. 5c, d). qPCR confirmed that genes related to the PI3K/AKT pathway were highly expressed in L-KO mice (Supplementary Fig. 4a). Similarly, p-AKT (S473 and T308) significantly decreased in human *PCK1*-OE cells but increased in *PCK1*-KO cells after 0.2 mM PA treatment (Fig. 5e, f).

**Fig. 5.**
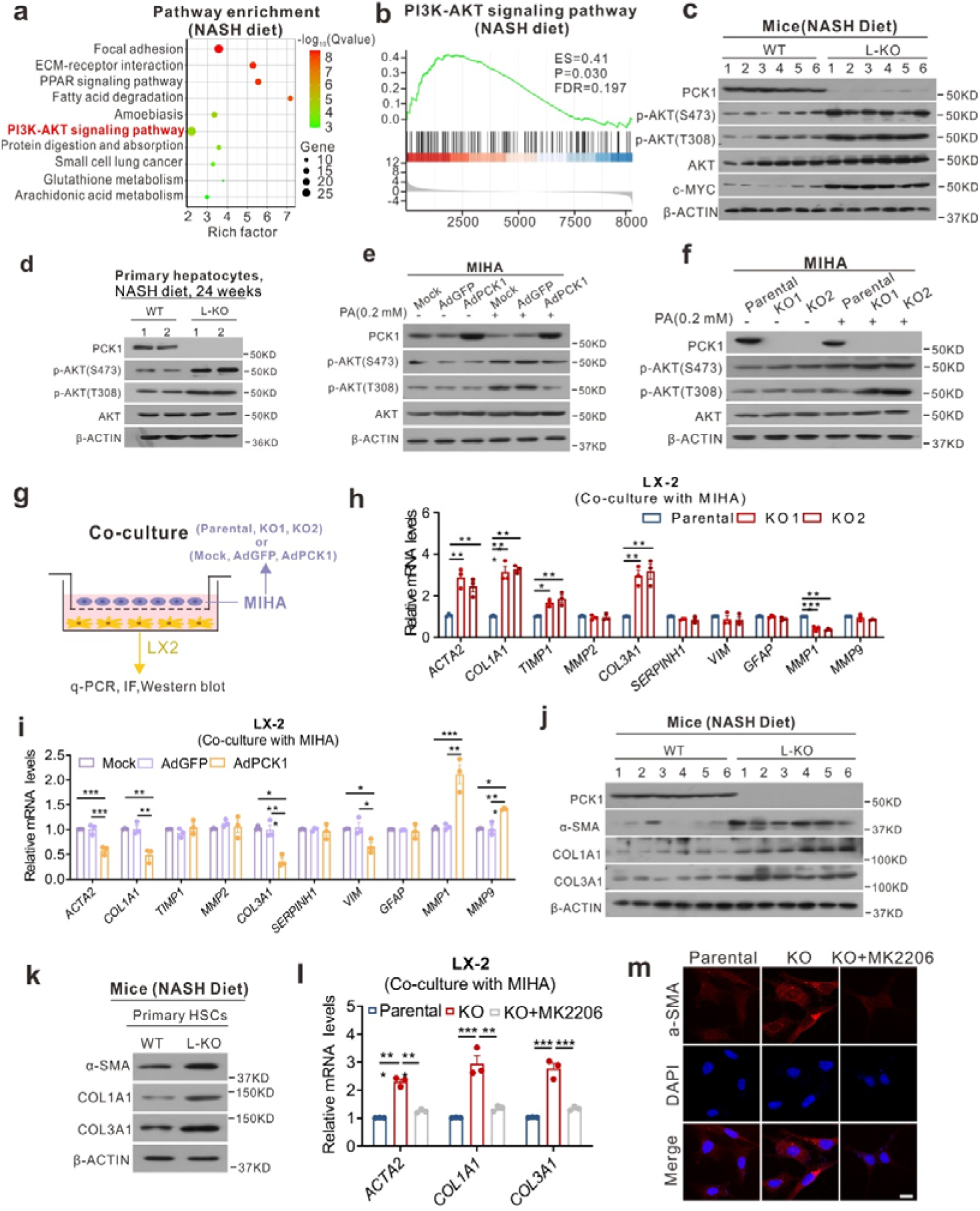
Hepatic PCK1 deficiency leads to HSC activation via PI3K/AKT pathway. **a** Pathway enrichment analysis of significantly upregulated genes in L-KO mice. **b** Gene Set Enrichment Analysis (GSEA) plot of enrichment in PI3K/AKT pathway. **c–f** Immunoblot analysis of AKT and p-AKT (S473 or T308) in mouse liver tissues (**c**), primary hepatocytes from NASH feeding mice(**d**), *PCK1*-OE (**e**), and *PCK1*-KO (**f**) MIHA cells with or without 0.2 mM palmitic acid (PA) treatment. **g** Schematic flow chart of co-culture models. **h, i** Quantitative PCR analysis of fibrosis-related genes in LX-2 cells co-cultured with *PCK1*-KO (**h**) or *PCK1*-OE (**i**) MIHA cells. **j, k** Western blotting of fibrosis related protein in liver tissues (**j**) or primary HSCs from NASH feeding mice (**k**) (n = 3) **l, m** Relative mRNA expression (**l**) and immunofluorescence images (**m**) of *ACTA2/*α-SMA, *COL1A1*, and *COL3A1* in LX-2 cells co-cultured with *PCK1*-KO MIHA cells treated with AKT inhibitor MK2206 (10 μM). Scale bars: 25 µm. Data are expressed as the mean ± SEM; \**p* < 0.05, \*\**p* < 0.01, \*\*\**p* < 0.001. *p* values obtained via two-tailed unpaired Student’s *t* tests or one-way ANOVA with Tukey’s post hoc test.

To clarify the role of PI3K/AKT pathway activation, transcriptome data were further analyzed. Interestingly, *Col1a1*, *Col3a1*, and *Lama2*, which are primary components of the extracellular matrix (ECM), were upregulated, as shown in the heat map of the PI3K/AKT pathway (Supplementary Fig. 4b). Moreover, GSEA analysis revealed that ECM-receptor interaction was upregulated in L-KO mice (Supplementary Fig. 4c). Because ECM deposition is typically considered as the key event underlying liver fibrosis, we predicted that the activation of the PI3K/AKT pathway promotes fibrosis in L-KO mice. HSCs are major ECM secretors; thus, we performed co-culture assays with human hepatocyte (MIHA) and human hepatic stellate cell lines (LX-2) (Fig. 5g). Interestingly, the mRNA levels of *ACTA2* (α-SMA, an HSC activation marker), *COL1A1*, and *COL3A1* increased in LX-2 cells co-cultured with *PCK1*-KO cells but decreased in LX-2 cells co-cultured with *PCK1*-OE cells (Fig. 5h, i). Similarly, COL1A1, COL3A1, and α-SMA expression increased in the liver tissues and primary HSCs of L-KO mice (Fig. 5j, k), which was confirmed in IHC analysis of COL3A1 (Supplementary Fig. 4d). However, these increases were partially reversed by MK2206, an AKT inhibitor (Fig. 5l, m). Collectively, these data suggest that the loss of PCK1 in hepatocytes induces HSC activation and ECM formation by activating the PI3K/AKT pathway.

### Paracrine PDGF-AA from hepatocytes promotes HSC activation

Hepatocytes elicit several fibrogenic actions in a paracrine manner to promote HSC activation ^20^. Thus, PCK1-mediated hepatic fibrosis may be involved in paracrine disorders. To test this hypothesis, several pro-fibrotic factors were screened; *Pdgfa* was found to be significantly elevated in the liver tissues of L-KO mice (Fig. 6a). Bioinformatics analysis confirmed that *PDGFA* was significantly increased in patients with NAFLD and NASH (Fig. 6b). *Pdgfa* encodes a dimer disulfide-linked polypeptide (PDGF-AA), and the chronic elevation of PDGF-AA in the mouse liver induces fibrosis ^21^. Immunoblotting and ELISA revealed increased PDGF-AA expression in the liver tissues, primary hepatocytes, and plasma of L-KO mice (Fig. 6c–f). Moreover, the PDGF-AA concentration markedly increased in the culture medium of *PCK1*-KO cells but decreased in that of *PCK1*-OE cells treated with 0.2 mM PA (Fig. 6g, h). Correspondingly, platelet-derived growth factor receptor alpha (*PDGFRA*), which encodes the PDGF-AA receptor, increased in LX-2 cells co-cultured with *PCK1*-KO cells but decreased in LX-2 cells co-cultured with *PCK1*-OE cells (Fig. 6i, j). To determine whether the pro-fibrogenic effect was mediated by PDGF-AA secretion, we treated the cells with a neutralizing antibody against PDGF-AA. As expected, the increases in α-SMA, COL1A1, and COL3A1 in LX-2 cells co-cultured with *PCK1*-KO cells were reversed by anti-PDGF-AA treatment (Fig. 6k).

**Fig. 6.**
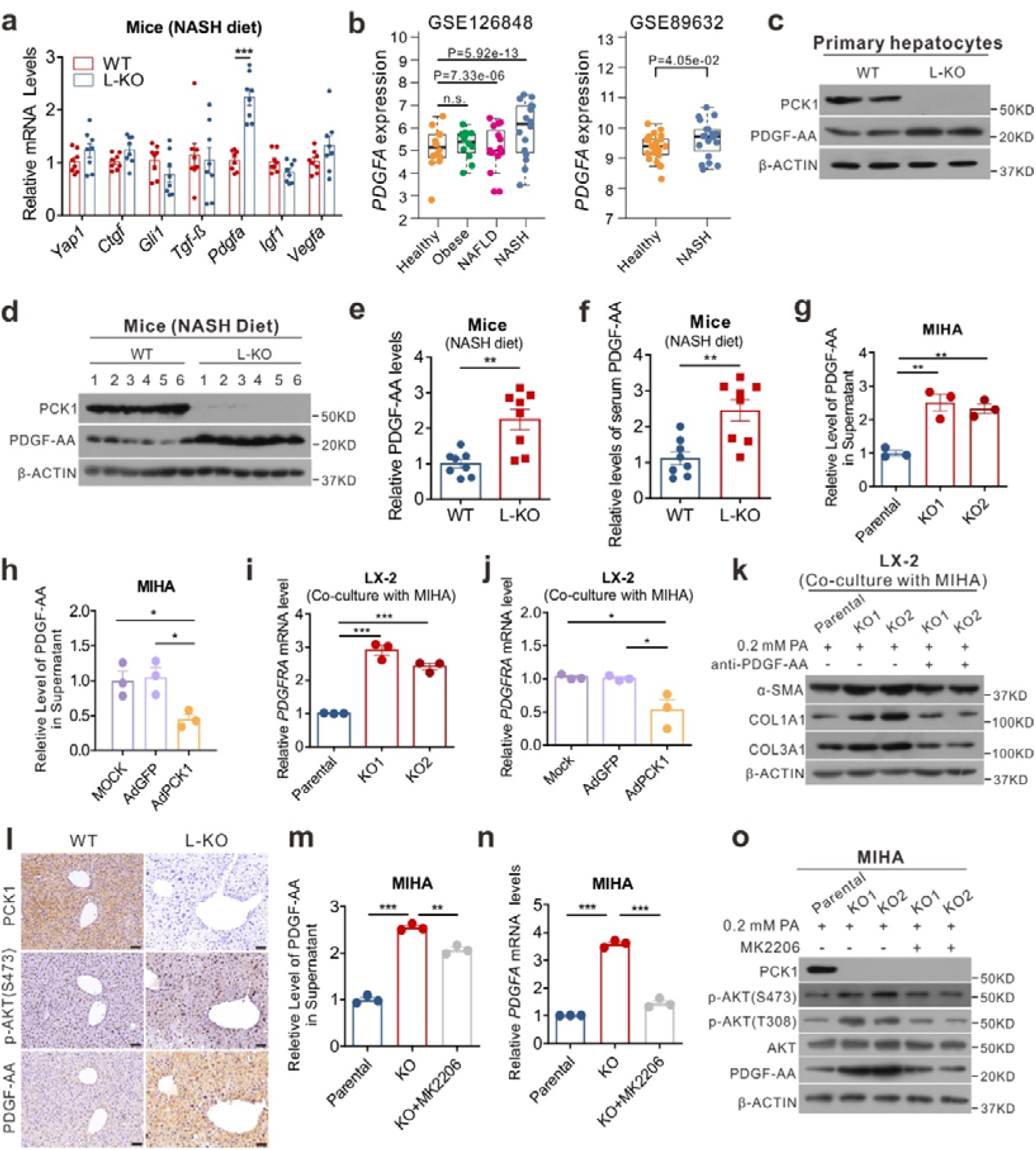
Paracrine PDGF-AA from hepatocytes promotes hepatic stellate cell (HSC) activation. **a** Expression levels of genes related to fibrogenesis. **b** Relative *PDGFA* mRNA levels in GSE126848 and GSE89632 datasets. **c, d** PDGF-AA protein levels in primary hepatocytes (**c**) or liver tissues (**d**) detected by western blotting. **e, f** PDGF-AA levels in liver tissues (**e**) or serum (**f**) were detected using enzyme-linked immunosorbent assay (ELISA). **g, h** Secreted PDGF-AA levels in conditioned medium with *PCK1*-KO (**g**) or *PCK1*-OE (**h**) MIHA cells treated with 0.2 mM palmitic acid (PA). **i**, **j** mRNA levels of *PDGFRA* in cell lysate of LX-2 cells co-cultured with *PCK1*-KO (**i**) or *PCK1*-OE (**j**) MIHA cells treated with PA. **k** Protein level in LX-2 cells co-cultured with *PCK1*-KO MIHA cells containing nonspecific rabbit IgG or a PDGF-AA blocking antibody. **I** Immunohistochemistry (IHC) analysis of PCK1, p-AKT (S473), and PDGF-AA in mouse liver sections (from serial sections). Scale bars: 50 µm. **m, n** Levels of PDGF-AA (**m**) or *PDGFA* (**n**) in conditioned medium or cell lysate of *PCK1*-KO MIHA cells treated with AKT inhibitor MK2206 (10 μM). **o** Protein levels in *PCK1*-KO MIHA cells treated with AKT inhibitor MK2206 (10 μM). Data are expressed as the mean ± SEM; \**p* < 0.05, \*\**p* < 0.01, \*\*\**p* < 0.001. *p* values obtained via two-tailed unpaired Student’s *t* tests or one-way ANOVA with Tukey’s post hoc test.

A review of the transcriptome data showed that *Pdgfa* appeared in the heat map of the PI3K/AKT pathway (Supplementary Fig. 4b). The IHC results showed that p-AKT (S473) was positively correlated with PDGF-AA (Fig. 6l). The AKT inhibitor MK2206 significantly blocked the increase in *PDGFA* levels in the supernatants or cells lysates of *PCK1*-KO cells (Fig. 6m–o). Taken together, these data confirm that PCK1 deficiency promoted PDGF-AA expression through the PI3K/AKT pathway and activated HSCs through hepatocyte-HSC crosstalk.

### PCK1 deficiency promotes the activation of the PI3K/AKT/PDGF-AA axis by activating RhoA signaling in hepatocytes

Rho GTPases, which cycle between active GTP-bound and inactive GDP-bound conformations, activate the PI3K/AKT pathway ^22–24^. Considering that PCK1 catalyzes the conversion of oxaloacetate to phosphoenolpyruvate, consuming GTP to generate GDP, we predicted that PCK1 deficiency alters intracellular GTP homeostasis. To test this hypothesis, high-performance liquid chromatography (HPLC) analysis was conducted to detect the intracellular levels of GTP. Interestingly, intracellular GTP levels decreased in *PCK1*-OE cells (Supplementary Fig. 5a) but increased in *PCK1*-KO cells (Supplementary Fig. 5b). Considering that Rho GTPases are activated when combined with GTP^25^, we examined the proteins levels of several Rho GTPases in the mouse liver tissues and found that GTP-bound RhoA significantly increased and inactivated RhoA, p-RhoA (S188), decreased in L-KO mice (Fig. 7a–c). Similar results were observed in the primary hepatocytes of mice fed the NASH diet (Supplementary Fig. 5c). Consistently, after PA treatment, the levels of GTP-bound RhoA decreased and p-RhoA (S188) expression increased in *PCK1*-OE cells, whereas the opposite results were observed in *PCK1*-KO cells (Fig. 7d–g). Next, RhoA inhibitor (Rhosin) or shRhoA was used to determine whether PI3K/AKT activation depends on RhoA. Immunoblotting, ELISA, and qPCR assays showed that both Rhosin and shRhoA blocked the increase in the activated forms of AKT and PDGF-AA in the *PCK1*-KO cell lysate and supernatant, as well as *ACTA2*, *COL1A1*, and *COL3A1* expression in LX-2 cells co-cultured with *PCK1*-KO hepatocytes (Fig. 7h–j, Supplementary Fig. 5d–f). To further evaluate the involvement of RhoA in HSC activation, we isolated primary hepatocytes from NASH-fed WT or L-KO mice; treated the cells with MK2206, Rhosin, or dimethyl sulfoxide vehicle; and then co-cultured the cells with primary HSCs from chow-fed WT mice. The activation of co-cultured HSCs was partially eliminated by the inhibition of AKT or RhoA in primary hepatocytes (Supplementary Fig. 5g). Moreover, PCK1 and p-RhoA (S188) were downregulated in samples from patients with NASH, whereas p-AKT (S473) and PDGF-AA levels were upregulated (Fig. 7k, l). These data indicate that PCK1 ablation stimulated the PI3K/AKT/PDGF-AA axis by activating RhoA.

**Fig. 7.**
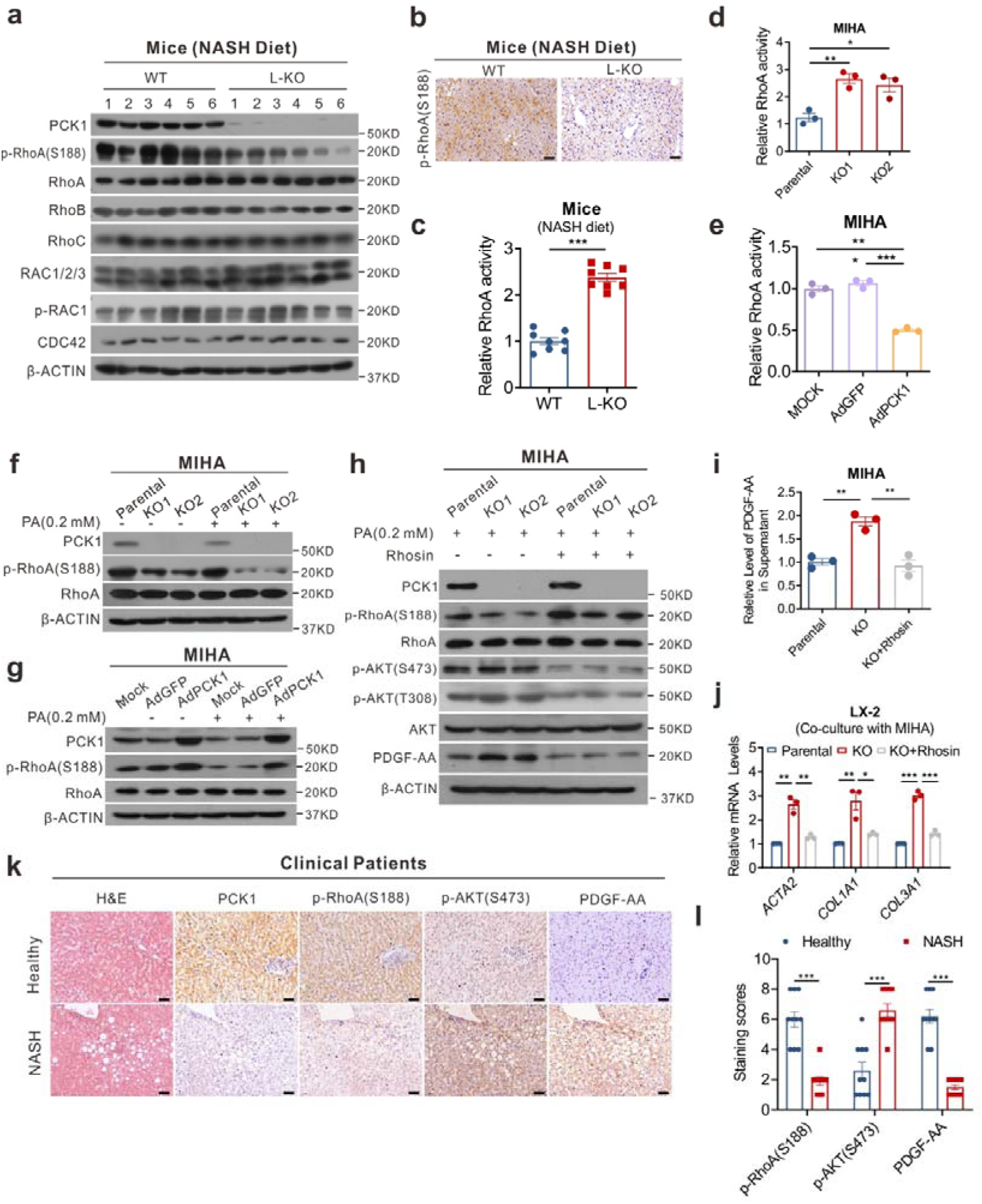
PCK1 deficiency promotes activation of PI3K/AKT/PDGF-AA axis by activating RhoA in hepatocytes. **a** Immunoblotting analysis of indicated proteins in mouse liver tissues. **b** Immunohistochemistry (IHC) analysis of p-RhoA (S188) in mouse liver tissues. Scale bars: 50 µm. **c–e** Relative levels of active RhoA were measured using G-LISA colorimetric RhoA activation assay in mouse liver tissues (**c**), *PCK1*-OE (**d**) and *PCK1*-KO (**e**) MIHA cells treated with 0.2 mM palmitic acid (PA). **f, g** Immunoblots of p-RhoA (S188) and RhoA in *PCK1*-OE (**f**) and *PCK1*-KO (**g**) MIHA cells with or without 0.2 mM PA treatment. **h** Expression of indicated proteins in *PCK1*-KO MIHA cells after addition of Rhosin (30 µM). **i** Levels of PDGF-AA in the supernatant of *PCK1*-KO MIHA cells treated with Rhosin (30 µM). **j** Relative mRNA expression of *ACTA2*, *COL1A1*, and *COL3A1* in LX-2 cells co-cultured with *PCK1*-KO MIHA cells treated with Rhosin (30 µM). **k** IHC analysis of PCK1, p-RhoA (S188), p-AKT (S473), and PDGF-AA in normal individuals and patients with NASH (from serial sections). Scale bars: 50 µm. **l** Semi-quantitative analyses of immunohistochemistry data of healthy and NASH human tissues for indicated proteins (healthy = 10, NASH = 10). Data are expressed as the mean ± SEM; \**p* < 0.05, \*\**p* < 0.01, \*\*\**p* < 0.001. *p* values obtained via one-way ANOVA with Tukey’s post hoc test.

### Genetic or pharmacological disruption of RhoA and AKT1 reduced progressive liver fibrosis *in vivo*

To explore whether blocking RhoA/PI3K/AKT could rescue the NASH phenotype in L-KO mice, the genetic and pharmacological disruption of RhoA and AKT1 was performed *in vivo* (Fig. 8a). Pharmacological inhibition of AKT1 or RhoA led to improved glucose intolerance (Supplementary Fig. 6a) and insulin resistance (Supplementary Fig. 6b). The increase in liver weight was also prevented (Fig. 8b), whereas the body weight decreased only in the MK2206 treatment group (Supplementary Fig. 6c). Additionally, Rhosin or MK2206 administration attenuated AST and ALT levels, as well as TG and FFA levels, in the serum and liver tissues (Figs. 8c, Supplementary Fig. 6d, e). Similarly, histochemistry revealed reduced liver steatosis, inflammation, and fibrosis in Rhosin- or MK2206-treated mice (Figs. 8d, Supplementary Fig. 6f), which was confirmed based on the decreased liver TNF-α and IL-6 levels (Fig. 8e). Additionally, α-SMA, COL1A1, COL3A1, PDGF-AA, and p-AKT (S473, T308) expression and GTP-bound RhoA levels decreased, whereas p-RhoA (S188) expression increased in the treatment group (Supplementary Fig. 6g–i). MK2206 or Rhosin treatment also reduced the expression of genes related to inflammation and fibrosis (Supplementary Fig. 6h). These results indicate that the pharmacological inhibition of AKT or RhoA can alleviate the clinical phenotypes of NASH.

**Fig. 8.**
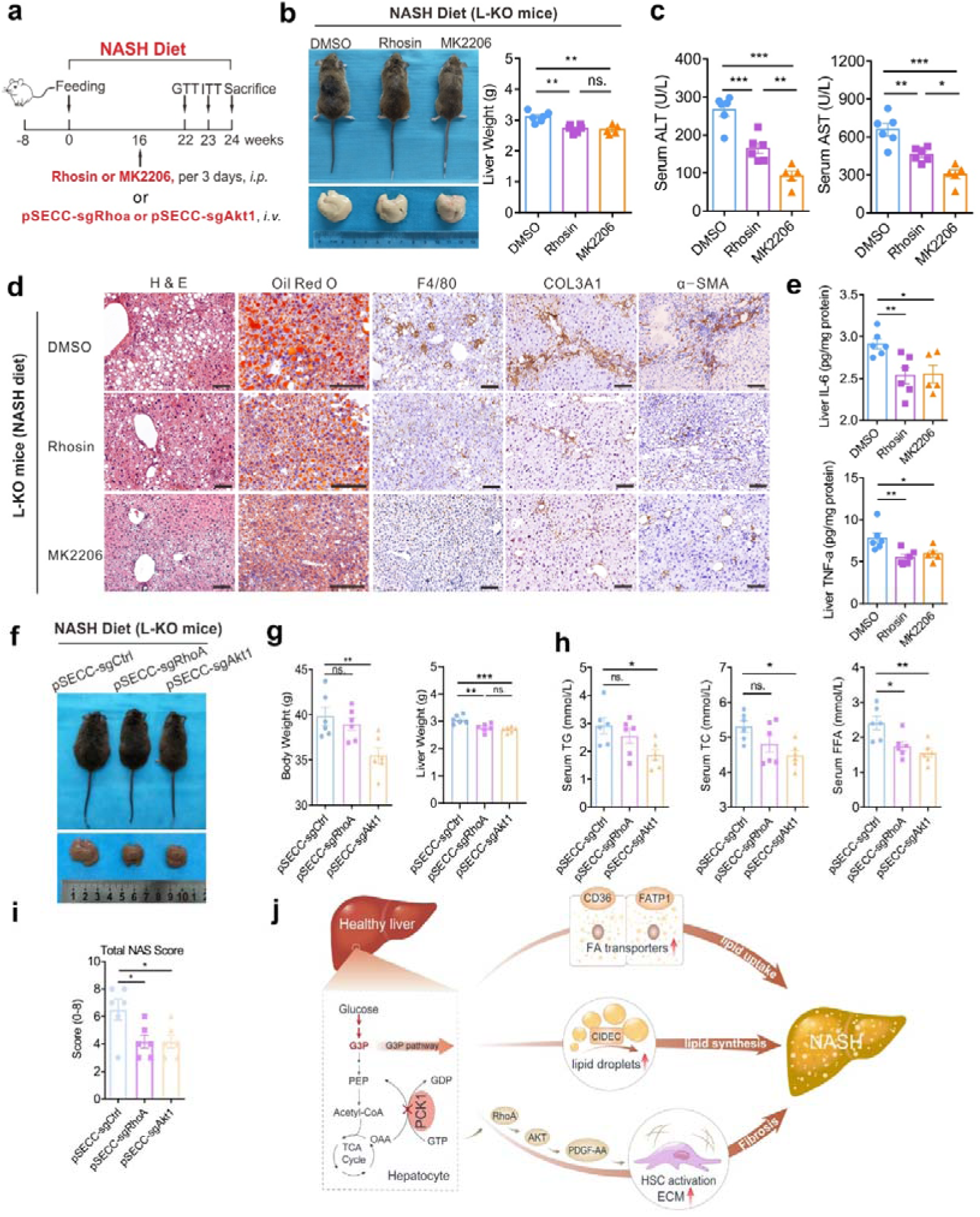
Pharmacological inhibition or genetic silencing of AKT1 or RhoA prevents NASH development *in vivo.* L-KO mice were fed the NASH diet for 24 weeks, and therapeutic treatments were initiated at different times. **a** Schematic diagram of *in vivo* pharmacological inhibition (**b**–**e**) and pSECC lentivirus-mediated silencing (**f**–**i**) of AKT1 or RhoA. **b, c** Representative whole-body, gross liver morphology, liver weight (**b**), and serum alanine aminotransferase (ALT) and aspartate aminotransferase (AST) (**c**). **d** Paraffin-embedded liver sections were stained with hematoxylin and eosin, or immunostained for F4/80, COL3A1, and α-SMA. Frozen sections stained with Oil Red O. Scale bars: 50 µm. **e** Levels of TNF-α and IL-6 in liver tissues. **f** Representative gross images of the livers from different groups of mice. **g** Quantification of body weight and liver weight in pSECC-sgAkt1 and pSECC-sgRhoA L-KO mice. **h** Plasma levels of total triglycerides (TG), total cholesterol (TC), and free fatty acids (FFA) were measured. **i** Normalized activity scores of liver sections. **j** Model depicting the critical role of PCK1 in controlling NASH progression. Data are expressed as the mean ± SEM; \**p* < 0.05, \*\**p* < 0.01, \*\*\**p* < 0.001; n.s., not significant. *p* values obtained via one-way ANOVA with Tukey’s post hoc test.

Next, we used pSECC, a lentiviral-based system that combines the CRISPR system and Cre recombinase, to silence RhoA or AKT1 in L-KO mice ^26^. In agreement with the results of the experiments described above, AKT or RhoA depletion *in vivo* partially prevented and liver weight and body weight gain (Fig. 8f,g). Moreover, serum AST, ALT, TC, TG, and FFA levels significantly decreased in sgAkt1- or sgRhoA-treated mice compared with those in sgCtrl-treated L-KO mice (Fig. 8h, Supplementary Fig. 7a). Histological analysis revealed that the NAS score, hepatic fibrosis, and F4/80^+^ macrophage counts decreased in sgAkt1 and sgRhoA mice, indicating the attenuation of the NASH phenotypes (Fig. 8i, Supplementary Fig. 7b, c). Consistently, genes or proteins involved in hepatic inflammation and fibrosis were significantly downregulated in the livers of sgAkt1 and sgRhoA mice, accompanied by the reduced expression of PDGF-AA (Supplementary Fig. 7d, e). These results indicate that the genetic inhibition of AKT or RhoA can alleviate the clinical phenotypes of NASH.

## Discussion

We found that the hepatic gluconeogenic enzyme PCK1 plays an important role in NASH progression. The expression of PCK1 was diminished in the livers of patients or mice with NASH. Moreover, deletion of PCK1 significantly exacerbated hepatic steatosis, fibrosis, and inflammation in mouse models fed the NASH diet. Mechanistically, loss of PCK1 not only promotes steatosis by enhancing lipid deposition, but also induces fibrosis through HSC activation via paracrine secretion of PDGF-AA, thus promoting NASH progression (Fig. 8j).

Abnormal lipid metabolism is characteristic of NAFLD and NASH. Previous studies predicted that altered lipid homeostasis was caused by abnormal expression of genes related to lipid metabolism ^27^. However, recent studies demonstrated that disruption of gluconeogenesis also leads to abnormal lipid metabolism. A deficiency of fructose-1,6-bisphosphatase 1 and glucose-6-phosphatase catalytic subunit, which are key enzymes in gluconeogenesis, results in severe hepatic steatosis and hypoglycemia, indicating that the suppression of gluconeogenesis also disrupts lipid homeostasis ^16,28^. As the first rate-limiting enzyme in gluconeogenesis, it is currently unclear whether PCK1 plays a critical role in NAFLD/NASH development. We identified a robust decrease in PCK1 expression in the livers of NASH mice and patients with NAFLD/NASH, causing severe hepatic steatosis and confirming that disordered hepatic gluconeogenesis affects lipid homeostasis.

Previous reports showed that PCK1 expression is increased in several obesity/diabetes mouse models, such as ZDF rats and *ob/ob* and *db/db* mice, and the disease progression of NASH is positively correlated with obesity and type 2 diabetes mellitus ^29–31^. Interestingly, we found that PCK1 expression was downregulated in a diet-induced murine NASH model. This discrepancy may be related to differences in the animal models used in different studies. Widely used rodent models for genetic forms of obesity and diabetes, such as *ob/ob* and *db/db* mice, exhibit increased plasma glucocorticoids, which may drive PCK1 expression ^31,32^. Another explanation is that the high-fat diet supplemented with high fructose/glucose in drinking water could have suppress PCK1 expression ^33^. Under a high-fat diet, PA can decrease the expression of PCK1 by inhibiting the expression of SIRT3 ^34^. Additionally, acetylation, ubiquitination, and phosphorylation modulate PCK1 expression ^35^. A high glucose level can reportedly destabilize PCK1 by stimulating its acetylation, thus promoting its ubiquitination and subsequent degradation ^36^. In this study, we found that ATF3, a member of the basic leucine zipper family of transcription factors ^37^, transcriptionally repressed *PCK1* upon PA overload *in vitro* and in the NASH mouse model. This result agrees with those of previous studies suggesting that ATF3 is upregulated in patients with NAFLD and murine NASH models and inhibits the expression of PCK1 in alcoholic fatty liver disease ^16,38,39^. Therefore, in this study, we found that PCK1 markedly decreased in NASH, and PA inhibited *PCK1* transcription by upregulating ATF3.

Numerous studies using PCK1 agonists or whole-body *Pck1* knockdown mice have verified that PCK1 can affect lipid metabolism ^40,41^. In the present study, liver-specific *Pck1* knockout induced significant hepatic steatosis even under normal feeding conditions. This is very important because it is uncommon for single gene ablation to cause spontaneous steatosis unless a high-fat diet is used. Moreover, mice with liver *Pck1* deficiency present aggravated inflammation when fed a high-fat high-fructose diet, contrasting with the results of a previous study showing that whole-body *Pck1* knockdown prevents hepatic inflammation ^42^. This discrepancy may be related to differences between diets and animal models, as whole-body *Pck1* knockdown may have unexpected effects on glucolipid metabolism. However, whether the restoration of PCK1 expression in the NASH model of L-KO mice can reverse NASH progression requires further investigation.

Lipid accumulation is characteristic of steatosis. Emerging evidence indicates that increased fatty acid uptake is associated with lipid accumulation ^41,42^. Previous studies demonstrated that the loss of PCK1 in the liver disturbed hepatic cataplerosis and led to the accumulation of TCA cycle intermediates. The slowed TCA cycle impaired fatty acid oxidation, resulting in fat accumulation in the liver ^43^. Additionally, elevated plasma TGs and FFAs could contribute to body weight gain in L-KO mice. Notably, NASH diet-induced glucose intolerance and insulin resistance in L-KO mice play essential roles in obesity and whole-body metabolism. In this study, genes involved in fatty acid uptake such as *Cd36* and *Slc27a1* were highly expressed in L-KO mice. In addition, a lipid droplet-associated protein Cidec was increased by both the chow and NASH diets, and was shown to be upregulated in patients with NALFD and L-KO mice, suggesting that PCK1 ablation promotes lipid droplet formation ^44,45^. Abnormal levels of metabolites also contribute to TG accumulation in the liver, with the G3P pathway contributing to over 90% of TG synthesis ^46^. As our metabolomics data showed that G3P and PA were significantly upregulated in L-KO mice, PCK1 deficiency may promote hepatic lipid accumulation by enhancing the expression of Cd36, Slc27a1, and Cidec and the levels of metabolic substrates such as G3P and PA. However, the precise mechanism by which PCK1 regulates the G3P pathway and expression levels of *Cd36* and *Slc27a1* must be further analyzed.

Fibrosis is another characteristic of NASH and drives the transition from simple steatosis to NASH. Activation of HSCs through the secretion of profibrotic cytokines, such as TGF-β and PDGF, is a key event in liver fibrosis ^43^. A recent study identified high mobility group protein B1, secreted by fructose-1,6-bisphosphatase 1-deficient hepatocytes, as the main mediator activating HSCs, revealing important crosstalk between hepatocytes and HSCs via paracrine signaling. Herein, PDGF-AA was secreted by PCK1-deficient hepatocytes and acted in a paracrine manner to activate HSCs. Increased deposition of ECM and activation of HSCs were observed in PDGFA-transgenic mice; however, the mechanism mediating PDGF-AA upregulation in fibrosis remains unclear ^21^. Here, we demonstrated that PCK1 deficiency promoted PDGF-AA secretion by activating the RhoA/PI3K/AKT pathway. Mechanistically, PCK1 deletion may increase intracellular GTP levels, thus promoting the activation of RhoA and further activating the PI3K/AKT pathway.

Most Rho GTPases cycle between an active GTP-bound and an inactive GDP-bound form, a process that is regulated by guanine nucleotide exchange factors, GTPase-activating proteins, and guanine nucleotide dissociation inhibitors. Guanine nucleotide exchange factors can activate Rho GTPases by catalyzing the exchange of GDP for GTP when the intracellular concentration of GTP is high ^44^. Several members of the Rho-GTPase family, such as Rac1, RhoA, and RhoC, can be activated by increased concentrations of intracellular GTP ^45–47^ ^48^. We found that PCK1 deficiency activated RhoA by increasing the levels of intracellular GTP, therefore activating the downstream PI3K/AKT pathway. Moreover, the genetic and pharmacological disruption of RhoA and AKT1 can effectively mitigate NASH phenotypes in L-KO NASH mice. In addition, RhoA and AKT inhibitors can reportedly inhibit the progression of NASH through other pathways, such as the NF-κB ^49^, Hippo ^50,51^, and Notch ^52^ signaling pathways. Although RhoA and AKT inhibitors are currently only in phase 3 trials or preclinical studies for the treatment of liver fibrosis or clinical tumors, these compounds also show therapeutic potential for NASH ^53–55^.

In conclusion, hepatic PCK1 deficiency promoted lipid deposition and fibrosis in a murine NASH model. Moreover, hepatic PCK1 loss activated the RhoA/PI3K/AKT pathway, which increased PDGF-AA secretion and promoted HSC activation. AKT/RhoA inhibitors reduced progressive liver fibrosis, providing a therapeutic strategy for NASH treatment.

## Materials and Methods

### Animal models

*Pck1*^loxp/loxp^ mice on a 129S6/SvEv background were purchased from the Mutant Mouse Resource & Research Center (MMRRC: 011950-UNC; Bar Harbor, ME, USA) and *Alb-Cre* mice on a C57BL/6 background were purchased from Model Animal Research Center of Nanjing University (Nanjing, China). To generate liver-specific *Pck1*-knockout mice (L-KO), *Alb-Cre* mice were crossed with *Pck1*^loxp/loxp^ mice. *Pck1*^loxp/loxp^ mice from the same breeding step were used as controls (wild-type, WT). Male WT and L-KO mice at 7–9 weeks old were fed the NASH diet (D12492: 60% Kcal fat, with drinking water containing 23.1 g/L fructose and 18.9 g/L glucose; Research Diets, New Brunswick, NJ, USA) (n = 11 per group) or control chow diet (D12450J: 10% Kcal fat, with tap water; Research Diets) (n = 10 per group) for 24 weeks. Food and drinking water were provided *ad libitum*. All mice were housed in temperature-controlled (23 °C) pathogen-free facilities with a 12 h light-dark cycle.

For AAV8 transduction, AAV8-TBG-*control* and AAV8-TBG-*Pck1* were purchased from the Shanghai Genechem Co., Ltd. (Shanghai, China) and injected via the tail vein following 10 weeks of NASH diet feeding (2 × 10^11^ genome copies/mouse). The efficiency of virus infection and expression of PCK1 in mouse hepatocytes were confirmed by western blot analysis. After a total of 24 weeks of NASH diet feeding, the mice were sacrificed for analysis. Genetic and pharmacological inhibition of AKT1 or RhoA were performed *in vivo*. Genetic depletion of mouse RhoA or AKT1 *in vivo* was conducted using pSECC (#60820; Addgene, Watertown, MA, USA), a lentiviral-based system that combined both the CRISPR system and Cre recombinase. After NASH diet feeding for 16 weeks, male L-KO mice were injected with pSECC-sgCtrl, pSECC-sgAkt1, or pSECC-sgRhoA through the tail vein at 1 × 10^9^ genome copies per mouse, with two booster injections at 7 day intervals. The pharmacological inhibition of AKT1 or RhoA was conducted using MK2206 or Rhosin. After NASH diet feeding for 16 weeks, the mice were divided into 3 groups and intraperitoneally injected with vehicle solution (n = 6), MK2206 (AKT inhibitor, 50 mg/kg, every 3 days) (n = 5), or Rhosin (RhoA inhibitor, 20 mg/kg, every 3 days) (n = 6) for 8 weeks. All mice were sacrificed for further study after NASH diet feeding for 24 weeks. Animal experiments were approved by the Animal Experimentation Ethics Committees of Chongqing Medical University and performed in accordance with the Guide for the Care and Use of Laboratory Animals.

### Liver tissues from patients with NASH

Paraffin-embedded normal (n = 10) and NASH human liver samples (n = 36) were kindly provided by Dr. Jiangao Fan, Dr. Xiaojun Wang, and Dr. Yalan Wang. Exclusion criteria were the presence of other causes of liver disease, including alcoholic fatty liver disease (>30 g/day for men, >20 g/day for women), chronic infection with hepatitis B and/or C virus, primary biliary cirrhosis, haemochromatosis, auto-immune hepatitis, and Wilson’s disease, as well as the use of anti-obesity, glucose-lowering, and/or lipid lowering pharmacological treatments. Liver tissue collection was approved by the Institutional Ethics Committees of Chongqing Medical University and Xin Hua Hospital. Informed consent was obtained from all subjects. The general characteristics of the NASH human liver samples are listed in **Supplementary Table 1**.

### Cell culture and treatment

All cell lines were grown in Dulbecco’s modified eagle medium (DMEM) supplemented with 10% fetal bovine serum, 100 µg/mL of streptomycin, and 100 U/mL of penicillin at 37 °C in 5% CO_2_. All cells were negative for mycoplasma. Short tandem repeat tests were performed to ensure the authenticity of the cells. Bovine serum albumin (BSA,10%), different concentrations of palmitic acid (PA), 10 µM MK2206 (AKT inhibitor), 40 µM Rhosin (RhoA inhibitor), or blocking antibody against PDGF-AA (2 µg/mL) was added to the medium. For *in vitro* co-culture assays, the human hepatic stellate cell (HSC) line LX-2 and human hepatocyte line MIHA cells (provided by Dr Ben C.B. Ko, The Hong Kong Polytechnic University, Hongkong, China) were used. LX-2 was pre-cultured in the lower chamber for 12 hours, and PCK1-OE or PCK1-KO cells (with or without indicated treatment) were seeded in Transwell inserts (#3401, Corning, NY, USA) that were subsequently loaded into the LX2-containing wells. The cells and supernatants were harvested for further analysis after 48 h of co-culture.

### Isolation of primary mouse hepatocytes

Primary hepatocytes were isolated and cultured from the livers of WT and L-KO mice fed the NASH diet as described previously ^56^. Briefly, following anesthesia, the inferior vena cava was cannulated and the liver was perfused *in situ* with 40 mL pre-warmed EGTA solution and 40 mL solution containing 0.35 mg/mL pronase (no. P5147, Sigma-Aldrich, St. Louis, MO, USA), followed by 40 mL solution containing 0.55 mg/mL collagenase (no. V900893, Sigma-Aldrich). After perfusion, the liver was crushed, and hepatocytes were released into the DMEM. The cell suspension was filtered through a 100 μm cell strainer and centrifuged at 50 ×*g* at 25°C for 3 min. After washing three times, the cells were suspended in DMEM supplemented with 10 mM glucose, 10% fetal bovine serum, 100 nM insulin (P3376, Beyotime Biotechnology, Shanghai, China), and 100 nM dexamethasone (D8040, Solarbio Life Sciences, Beijing, China), and then plated on 60 mm diameter plastic plates ^56^. After cell attachment, the medium was replaced with serum-free media, and the cells were used for experiments on the following day.

### Isolation of primary mouse hepatocytes and primary HSCs

HSCs were isolated from the mice as described previously ^57^. Briefly, after perfusion with solutions containing protease and collagenase, the crushed liver was digested with a solution containing 1% DNase (10104159001, Roche Diagnostics GmbH, Mannheim, Germany), 0.5 mg/mL protease, and 0.55 mg/mL collagenase for 25 min. The cell suspension was filtered through a 70 µm cell strainer, centrifuged at 580 ×*g* for 10 min at 4 °C, and washed twice with Gey’s balanced salt solution. The cells were subjected to gradient centrifugation on a 9.7% Nycodenz (1002424, Axis-Shield, Oslo, Norway) to isolate HSCs, which were then plated onto collagen-coated plates. The cells were cultured in DMEM containing 10% (vol/vol) fetal bovine serum and 1% penicillin-streptomycin ^58^.

### Construction of adenovirus, lentivirus, and stable cell lines

AdGFP and AdPCK1 adenoviruses were generated using the AdEasy system as described previously ^4^. The *PCK1* knockout (*PCK1*-KO) MIHA cell line was constructed using the CRISPR-Cas9 system (from Prof. Ding Xue, the School of Life Sciences, Tsinghua University, Beijing, China), as described previously ^4^. To knock down *ATF3* expression in MIHA cells, three pairs of oligonucleotides encoding short hairpin RNAs (shRNAs) targeting *ATF3* or negative control shRNA were cloned into the pLL3.7 vector (from Prof. Bing Sun, Shanghai Institute of Biochemistry and Cell Biology, Chinese Academy of Sciences, China). The lentiviruses were obtained by transient transfection of the psPAX2 packaging plasmid and pMD2.G envelope plasmid in HEK293T cells by using Lipofectamine 3000 Transfection Reagent (Thermo Fisher Scientific, Waltham, MA, USA). Transfection efficiency was validated by western blotting. Information on the reagents is listed in **Supplementary Table 2**. The pSECC lentiviral vector cloning and packaging strategy have been described previously^26^.

### RNA extraction and real-time PCR

Total RNA was extracted from the liver tissues or cell lines using Trizol reagent (15596018, Invitrogen, Carlsbad CA, USA) according to the manufacturer’s instructions. RNA was reverse-transcribed using a PrimeScript RT reagent Kit with gDNA Eraser (Cat: RR047A, TAKARA, Shiga, Japan), and qPCR was performed on a Bio-Rad CFX96 machine (Hercules, CA, USA). The mRNA levels of selected genes were calculated after normalization to β-actin by using the 2^(-ΔΔC(T))^ method. Primer sequences are provided in **Supplementary Table 3**.

### Immunoblotting

Mouse liver tissues and cells were homogenized and lysed in Protein Extraction Reagent (P0013, Beyotime Biotechnology) supplemented with protease inhibitor cocktail (04693159001, Roche, Basel, Switzerland). Equal amounts of protein lysates were separated by sodium dodecyl sulfate-polyacrylamide gel electrophoresis and transferred onto polyvinylidene fluoride membranes (Millipore, Billerica, MA, USA). The antibodies are listed in **Supplementary Table 4**. β-Actin was used as a loading control.

### BODIPY staining

The cells were fixed in 4% formaldehyde for 30 min at room temperature, permeabilized with 0.1% Triton X-100 for 5 min at room temperature, and stained with BODYPY (D3823, Invitrogen) for 60 min. Stained sections were analyzed using a Leica confocal microscope (Leica TCS SP8, Wetzlar, Germany).

### CCK8 assay

The cells were seeded at 5 × 10^3^ cells /well in 96-well plates and treated with different concentrations of PA for 24 h. 10 μL CCK-8 solution (#C0005, Topscience, Shanghai, China) was added to each well and incubated at 37 °C for 1 h. The absorbance was measured at 450 nm.

### ELISA and G-LISA

Serum insulin (P3376, Beyotime Biotechnology), TNFα (PT512, Beyotime Biotechnology), IL-6 (PI326, Beyotime Biotechnology), and PDGF-AA (SEA523Mu, Cloud-Clone Corp., Wuhan, China) concentrations in the liver tissue and plasma were quantified using mouse ELISA kits according to the manufacturer’s instructions. The levels of secreted PDGF-AA in the cell culture supernatants were determined using a human ELISA kit (SEA523Hu, Cloud-Clone Corp). Active RhoA was detected in a colorimetric RhoA activation assay (G-LISA) (BK124, Cytoskeleton, Denver, CO, USA).

### Immunofluorescence

LX-2 cells were fixed in 4% formaldehyde for 25 min and then incubated in 10% normal goat serum for 1 h. The cells were incubated with primary α-SMA antibodies. Specific signals were visualized using secondary antibodies (ZF-0316, Zsbio, Beijing, China). For nuclear staining, the cells were treated with 1 μg/mL DAPI (10236276001, Roche Diagnostics). The samples were detected with a laser-scanning confocal microscope (Leica TCS SP8).

### Chromatin immunoprecipitation (ChIP) assay

Chromatin immunoprecipitation (ChIP) and ChIP-quantitative real-time PCR (ChIP-qPCR) assays for MIHA cells were performed as described previously ^59^. Briefly, sonicated chromatin was used for the immunoprecipitation assay. The pre-cleared supernatants were incubated with a monoclonal antibody against ATF3 overnight at 4 °C, followed by a 4 h of incubation with protein A/G agarose beads (LOT: 3460992, Millipore). The purified DNA fragments bound by ATF3 were analyzed using qPCR. IgG and histone H3 were used as negative and positive controls, respectively. The primers used for real-time PCR analysis of PCK1 are listed in **Supplementary Table 3**.

### Glucose and insulin tolerance tests

The glucose tolerance test (GTT) and insulin tolerance test (ITT) were performed at 2 or 1 weeks prior to sacrifice. The animals were fasted for 16 or 6 h, and then glucose solution (2 g/kg body weight) or insulin (0.75 U/kg body weight) was administered via an intraperitoneal injection, respectively. Venous tail blood samples were collected at 0, 30, 60, 90, and 120 min post-administration to assess blood glucose levels using a glucose meter (HGM-114, OMRON, Kyoto, Japan).

### Biochemical analysis

The serum levels of aspartate transaminase (AST), alanine transaminase (ALT), total triglyceride (TG), total cholesterol (TC), and free fatty acids (FFA) were determined with an automated biochemical analyzer (Hitachi 7600, Tokyo, Japan). TG, TC, and FFA levels in the mouse liver tissues were measured with commercial kits according to the manufacturer’s protocol (TG: cat. no. BC0625; TC: cat. no. BC1985; FFA: cat. no. BC0595, Solarbio Life Sciences).

### Histological analysis

Paraffin blocks were sectioned into 4 μm slices and used for hematoxylin and eosin (HE) staining, Sirius red staining, and immunohistochemistry (IHC) assay according to standard protocols. Frozen liver tissue sections were stained with Oil Red O (G1260, Solarbio). For pathological grading, all liver specimens were scored by two experienced pathologists according to the NAFLD activity score (NAS), defined as the sum of steatosis (0–3), inflammation (0–3), and hepatocyte ballooning (0–2). An NAS score ≥5 was considered to indicate NASH. Samples were scanned using a slide scanner (Pannoramic DESK, 3D Histech kft, Hungary). For quantitative analysis, the areas of lipid droplets and Sirius red staining were quantified using ImageJ software (version 1.6.0; NIH, Bethesda, MD, USA). Immunohistochemical staining was semi-quantitatively analyzed using the immunoreactive scoring system ^60^. The percentage of positive cells was graded on a scale of 0−4: (0: negative, 1: 0–25%, 2: 26–50%, 3: 51–75%, 4: 76–100%). The signal intensity was scored on a scale of 0–3: 0 = negative; 1 = weak; 2 = moderate; and 3 = strong. Thus, the final immunoreactive score□=□(score of staining intensity) × (score of percentage of positive cells).

### Transcriptomic analyses

Using Trizol reagent, total RNA was isolated from the liver tissues of WT and L-KO mice fed a chow or NASH diet. The RNA quality was checked with a Bioanalyzer 2200 (Agilent Technologies, Santa Clara, CA, USA). cDNA libraries were prepared using an NEBNext^®^ Ultra™ Directional RNA Library Prep Kit, NEBNext^®^ Poly (A) mRNA Magnetic Isolation Module, NEBNext^®^ Multiplex Oligos according to the manufacturer’s instructions (New England Biolabs, Ipswich, MA, USA). Genes with fold-change >2.0 or <0.5 and false discovery rate <0.05 were considered to be significantly differentially expressed. The volcano, heat, and bubble maps were generated using the ‘ggplot2’ or ‘ggpubr’ packages in R (version 3.6.3; The R Project for Statistical Computing, Vienna, Austria). Gene set enrichment analysis was performed using ‘enrichplot’ packages ^61^. The RNA-seq data files have been deposited to the Gene Expression Omnibus database (www.ncbi.nlm.nih.gov/geo/) under accession number GSE162211.

### Untargeted metabolomics

Following sacrifice, the livers were snap-frozen in liquid nitrogen and stored at −80 °C until analysis. Untargeted metabolomics was performed using an ultra-high performance liquid chromatography apparatus (Agilent 1290 Infinity LC, Agilent Technologies) coupled to a quadrupole time-of-flight (TripleTOF 6600, AB SCIEX, Framingham, MA, USA) at Shanghai Applied Protein Technology Co., Ltd. (Shanghai, China). Metabolites with a variable importance in projection value >1 were evaluated using Student’s *t*-test. *p* < 0.05 was considered to indicate statistically significant results.

### HPLC anaIysis of celluIar nucIeotides

Cellular nucleotides were extracted according to published procedures ^62^. Briefly, 1 × 10^6^ cells were washed with phosphate-buffered saline and quenched with liquid nitrogen, and then vigorously mixed with methanol and acetonitrile (1:1, v:v). After incubation on ice for 15 min, the samples were centrifuged at 12,000 ×*g* at 4 °C for 10 min. Cellular nucleotides were separated and quantified using a C18 column (Agilent Eclipse XDB-C18, 4.6 × 250 mm, average particle size 5 μm) assembled on the Waters Alliance e2695 Separations Module (Milford, MA, USA). Acetonitrile (5%) and 50 mM KH_2_PO_4_ (pH 6.5) containing 10 mM tetrabutylammonium bromide were used as mobile phase A, and acetonitrile was used as mobile phase B. All samples were separated in the mobile phase at a flow rate of 1 mL/min for 30 min at 22 °C. No degradation of individual nucleotides or changes in the ratios of nucleotide mixtures was detected during the experiments. The GTP concentration in the samples was calculated based on the slope of the calibration curves generated using pooled authentic samples (to mimic the matrix), and guanosine-5′-triphosphoric acid disodium salt was used as a standard.

### GEO database mining

Raw data in the GSE126848, GSE89632, and GSE135251 datasets were downloaded from the GEO database (https://www.ncbi.nlm.nih.gov/geo/). R package DESeq2 was used to analyze gene expression levels between different samples, such as NASH vs. healthy, NAFLD vs. healthy, and obesity vs. healthy. Genes showing |log_2_ fold-change|>1 and false discovery rate <0.05 were considered to present differential expression.

### Statistics

Statistical analyses were performed using GraphPad Prism 6.0 software (GraphPad, Inc., La Jolla, CA, USA). Data were represented as the mean ± standard error of the mean unless otherwise stated. Significant differences between the means of two groups were determined using Student’s *t*-test. One-way analysis of variance was used to determine statistical significance for experiments with more than two groups followed by Tukey’s post hoc test. For correlation analysis, Spearman’s correlation coefficient was used. *p* < 0.05 was considered to indicate statistically significant results.

## Acknowledgements

We would like to thank Dr. T.-C He (University of Chicago, USA) and Prof. Ding Xue (Tsinghua University, China) for providing the pAdEasy and CRISPR/Cas9 system, respectively. We thank Prof. Youde Cao and Yalan Wang (Chongqing Medical University, China) for providing samples and pathological analysis support. This work was supported by the National Natural Science Foundation of China (grant no. U20A20392, 82073251, 82072286, 81872270), the 111 Project (No. D20028), the Natural Science Foundation Project of Chongqing (cstc2018jcyjAX0254, cstc2019jscx-dxwtBX0019), the Leading Talent Program of CQ CSTC (CSTCCXLJRC201719), the Science and Technology Research Program of Chongqing Municipal Education Commission (HZ2021006, KJZD-M202000401), the Future Medical Youth Innovation Team of Chongqing Medical University (W0036, W0101), the Kuanren talents program of the second affiliated hospital of Chongqing Medical University, the Scientific Research Innovation Project for Postgraduate in Chongqing (CYB19168, CYS19193), and the Youth talent fund of Sichuan Provincial People’s Hospital in 2020 (No. 2020QN03).

## Author contributions

NT, AH, and KW conceived and designed the study. QY, YL and GZ performed most experiments and analyzed the data. HD, CC and KW conducted bioinformatics analysis. XP assisted with mice experiments. XW, JF, and QP provided human NAFLD/NASH samples. QY, KW, and NT wrote the manuscript with all authors providing feedback. The order of the co-first authors was assigned on the basis of their relative contributions to the study.

## Ethical approval

Animal experiments were approved by Animal Experimentation Ethics Committees of Chongqing Medical University and were carried out in accordance with the Guide for the Care and Use of Laboratory Animals. Liver tissue collection was approved by the Institutional Ethics Committees of Chongqing Medical University and Xin Hua Hospital. Informed consent was obtained from all subjects.

## Data availability

RNA-seq data files have been deposited into the Gene Expression Omnibus database (www.ncbi.nlm.nih.gov/geo/) with accession number GSE162211. All other data are available from the corresponding author upon reasonable request. Source data are provided with this paper.

