## Supplementary for "Deficiency of gluconeogenic enzyme PCK1 promotes non-alcoholic steatohepatitis progression and fibrosis through PI3K/AKT/PDGF axis activation"

### Supplementary Figures and Figure Legends

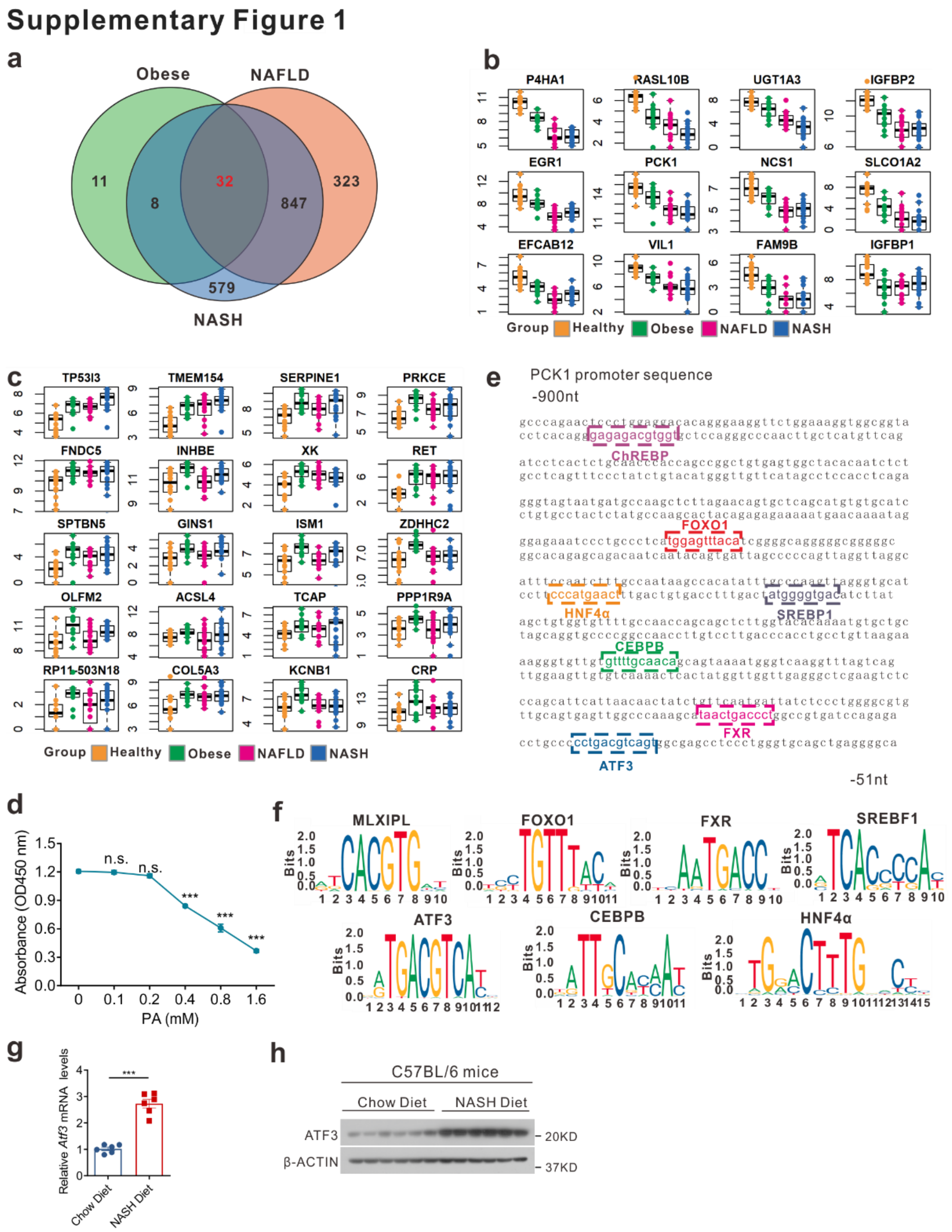


**Supplementary Fig. 1 PCK1 is downregulated in NASH patients due to ATF3 upregulation in NAFLD/NASH mouse model. a** Venn Diagram showing a total of 32 genes were significantly changed in patients with obesity, NAFLD, and NASH. **b** 12 genes were significantly downregulated in obesity, NAFLD, and NASH. **c** 20 genes were significantly upregulated in obesity, NAFLD and NASH. **d** Cell proliferation was assessed by a CCK8 assay. **e** Illustration of the predicted transcription factor binding sites in the 0.9 kb *PCK1* promoter region using JASPAR. **f** Potentially regulatory binding sequences of the transcription factors generated by JASPAR. **g, h** The mRNA (**g**) and protein levels (**h**) of ATF3 in mice fed with NASH diet. Data expressed as mean ± SEM; $^{*}P < 0.05$, $^{**}P < 0.01$, $^{***}P < 0.001$. *P* values obtained via 2-tailed unpaired Student's t tests or one-way ANOVA with Tukey's post hoc test.

### Supplementary Figure 2

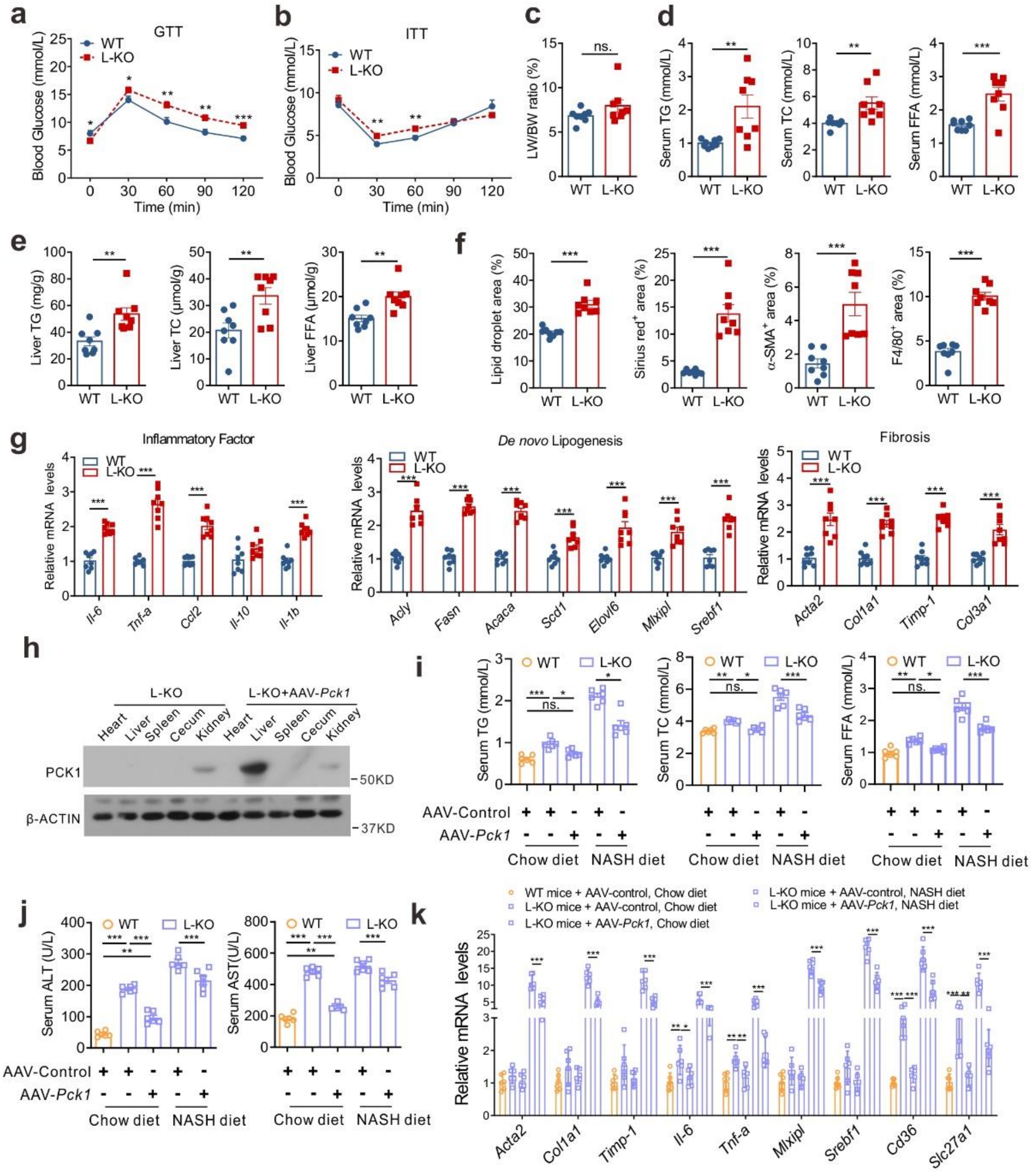


**Supplementary Fig. 2 L-KO mice had abnormal lipid metabolism and severe liver injury when fed NASH diet.** For **a-g**, WT and L-KO mice were treated with NASH diet for 24 weeks, n=8/group. For **h-j**, mice were administrated with AAV8-TBG-*Control* or AAV8-TBG-*Pck1* after 10 weeks of chow diet or NASH diet feeding, n=6/group. **a, b** GTT (**a**) and ITT (**b**) in WT and L-KO mice after 24 weeks of NASH diet. **c** Liver weight to body weight ratio of mice from the indicated groups. **d, e** TG, TC, and FFA levels in serum (**d**) or liver tissues (**e**). **f** Quantifications of Oil red O staining, Sirius red staining, and IHC staining. **g** Genes associated with inflammatory infiltration, *de novo* lipogenesis, and fibrogenesis were measured in WT and L-KO mice. **h** Western blot analysis of exogenous PCK1 protein levels in the livers of AAV–injected mice. **i-j** Levels of serum TG, TC, FFA, ALT and AST concentrations were measured. **k** qPCR analysis of genes associated with inflammatory infiltration, *de novo* lipogenesis and fibrogenesis in mice administrated with AAV8-TBG-Control or AAV8-TBG-*Pck1*. Data expressed as mean ± SEM; *$P < 0.05$, ** $P < 0.01$, ***$P < 0.001$. $P$ values obtained via 2-tailed unpaired Student's t tests.

### Supplementary Figure 3

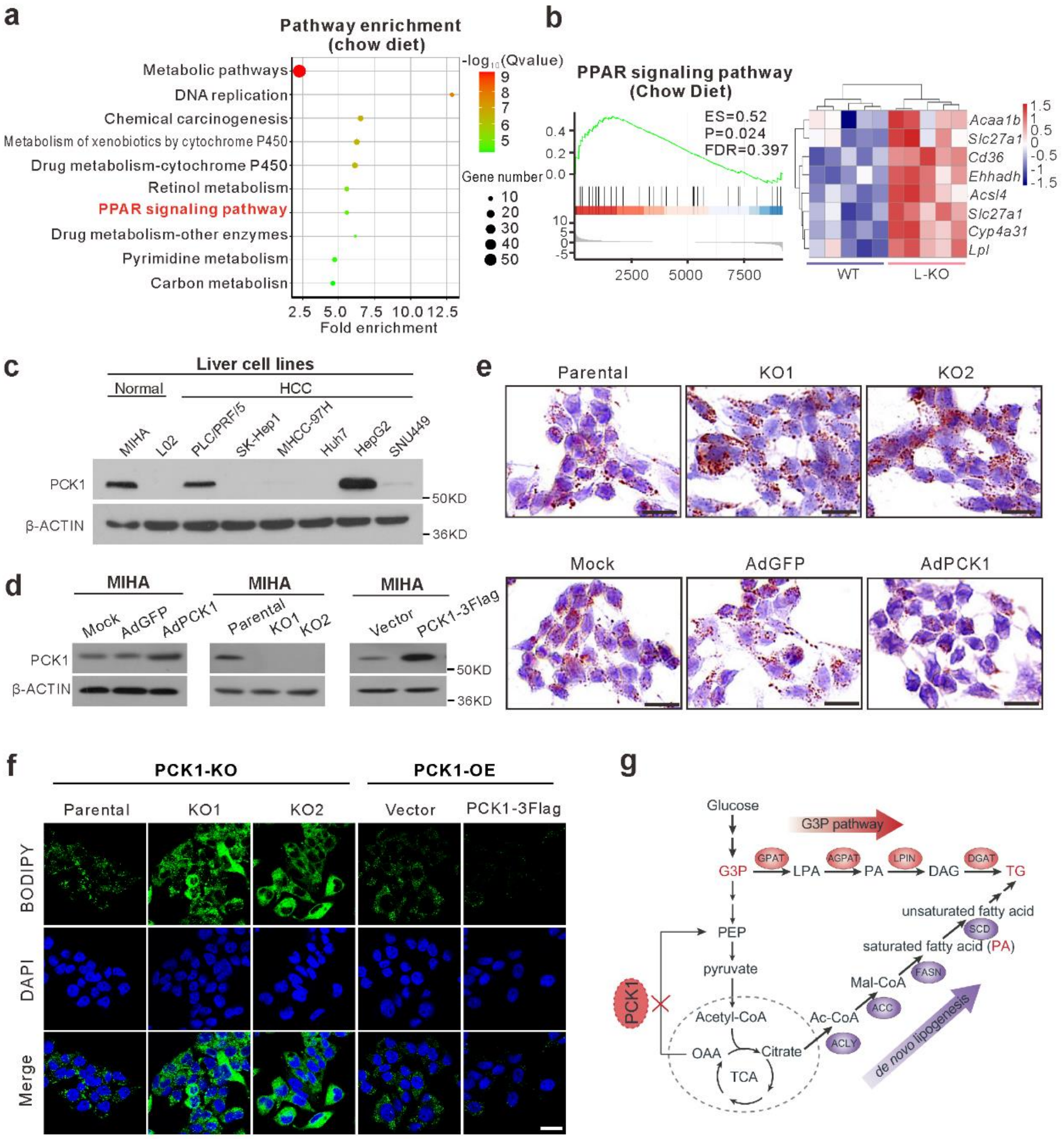


**Supplementary Fig. 3 Loss of PCK1 exacerbates lipid metabolism dysfunction *in vitro*. a** Pathway enrichment analysis of L-KO mice fed chow diet (n=5). **b** GSEA revealed that the transcription level of PPAR signaling pathway was prominently upregulated in L-KO mice fed chow diet (n=5). **c** Expression levels of PCK1 in different cell lines were measured by western blot. **d** Protein levels of PCK1 were detected by western blot analysis in *PCK1-OE*

and *PCK1-KO* MIHA cells. **e, f** Representative Oil Red O staining (**e**) and BODIPY staining (**f**) in *PCK1-OE* and *PCK1-KO* MIHA cells. Scale bars: 25 μm. **g** Schematic presentation of the G3P pathway and DNL. G3P: glycerol-3-phosphate; PA: palmitic acid; DL: de novo lipogenesis.

### Supplementary Figure 4

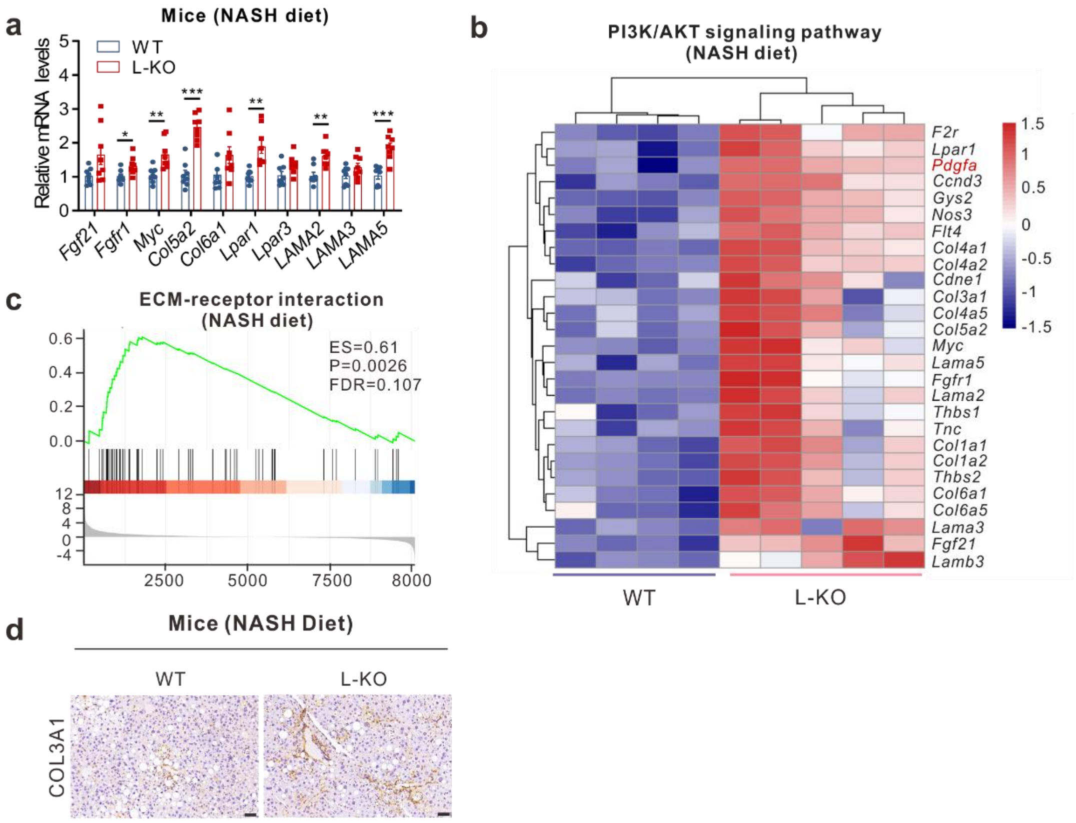


**Supplementary Fig. 4 PI3K/AKT signaling pathway and ECM-receptor interaction were activated in L-KO mice fed NASH diet. a** mRNA levels of the indicated genes in the liver of WT and L-KO mice (n=8). **b** Heat map of expression levels of PI3K/AKT target genes in WT and L-KO mice (n=4-5). **c** GSEA revealed the "ECM-receptor interaction" was upregulated in L-KO mice (n=4-5). **d** COL3A1 immunostaining in mice liver sections. Scale bars: 50 μm. Data expressed as mean ± SEM; *$P < 0.05$, ** $P < 0.01$, ***$P < 0.001$. *P* values obtained via 2-tailed unpaired Student's t tests.

### Supplementary Figure 5

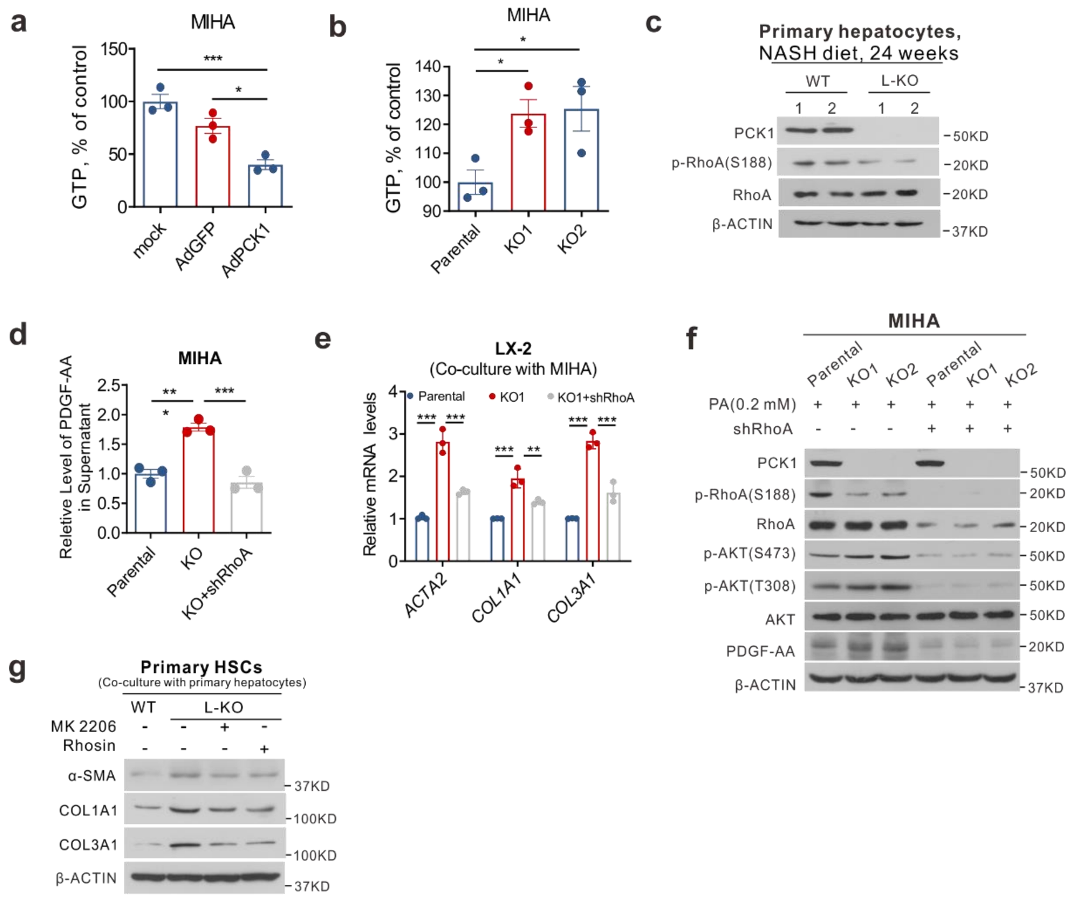


**Supplementary Fig. 5 PCK1 deficiency promoted the accumulation of GTP, and RhoA knockdown reversed the activation of PI3K/AKT/PDGF-AA axis.** **a-b** The intracellular GTP levels in *PCK1*-OE (**a**) and *PCK1*-KO (**b**) cells treated with 0.2 mM PA were determined by HPLC. **c** Immunoblots of the indicated proteins in the primary hepatocytes isolated from WT and L-KO mice fed NASH diet. **d** Levels of PDGF-AA in the supernatant of *PCK1*-KO MIHA cells infected with either shControl or shRhoA treated with 0.2 mM PA. **e** Relative mRNA expression of *ACTA2*, *COL1A1*, and *COL3A1* in LX-2 cells co-cultured with *PCK1*-KO MIHA cells infected with either shControl or shRhoA. **f** Immunoblot analysis of indicated proteins in *PCK1*-KO MIHA cells infected with either shControl or shRhoA. **g** Isolated mouse primary hepatocytes from WT and L-KO mice fed NASH diet for 24 weeks were treated with MK2206, Rhosin or DMSO vehicle, and then co-cultured with primary hepatic stellate cell isolated from WT mice fed chow diet, and the protein levels were determined. Data expressed as mean ± SEM; **P* < 0.05, ** *P* < 0.01, ****P* < 0.001. *P* values obtained via 2-tailed unpaired Student's t tests.

### Supplementary Figure 6

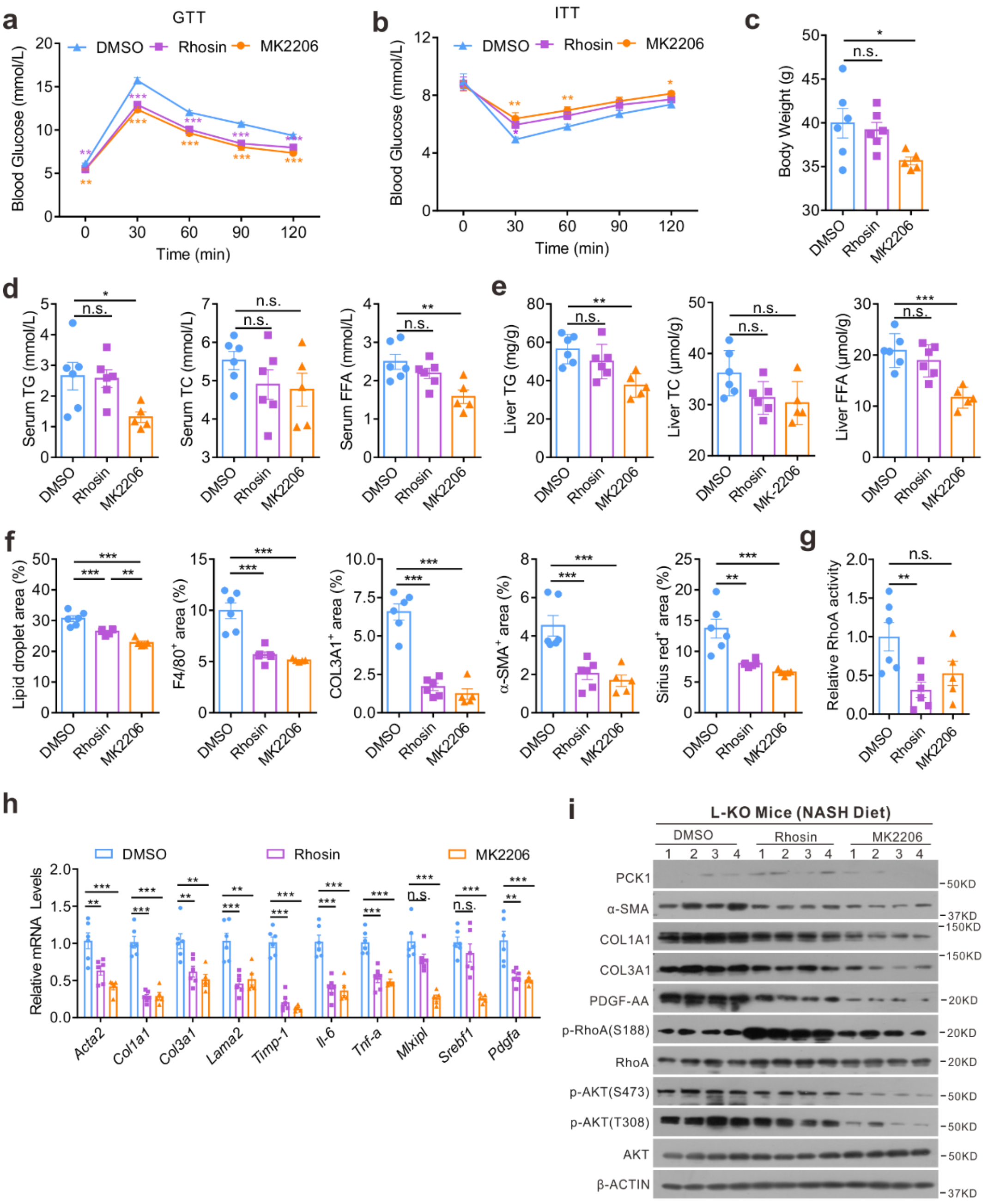


**Supplementary Fig. 6 RhoA/AKT inhibition partially relieves NASH phenotype in L-KO mice fed NASH diet. a, b** Glucose levels measured during the glucose tolerance test (GTT) (**a**) and insulin tolerance test (ITT) (**b**). **c** Body weight of mice in the indicated groups. **d** Serum TG, TC, and FFA levels were measured using automated biochemical analyzer. **e** Liver TG, TC, and FFA levels determined by colorimetry. **f** Quantification of liver sections of L-KO mice treated with DMSO, Rhosin or MK2206. **g** Reletive GTP-bound RhoA levels in mice liver tissues. **h** mRNA levels of genes associated with lipid metabolism, fibrogenesis, and inflammatory infiltration. DMSO group (n=6), Rhosin group (n=6), MK2206 group (n=5). **i** Expression of the indicated proteins in mice liver tissues. Data expressed as mean ± SEM; **P* < 0.05, ** *P* < 0.01, ****P* < 0.001; n.s., not statistically significant. *P* values obtained via one-way ANOVA with Tukey's post hoc test.

### Supplementary Figure 7

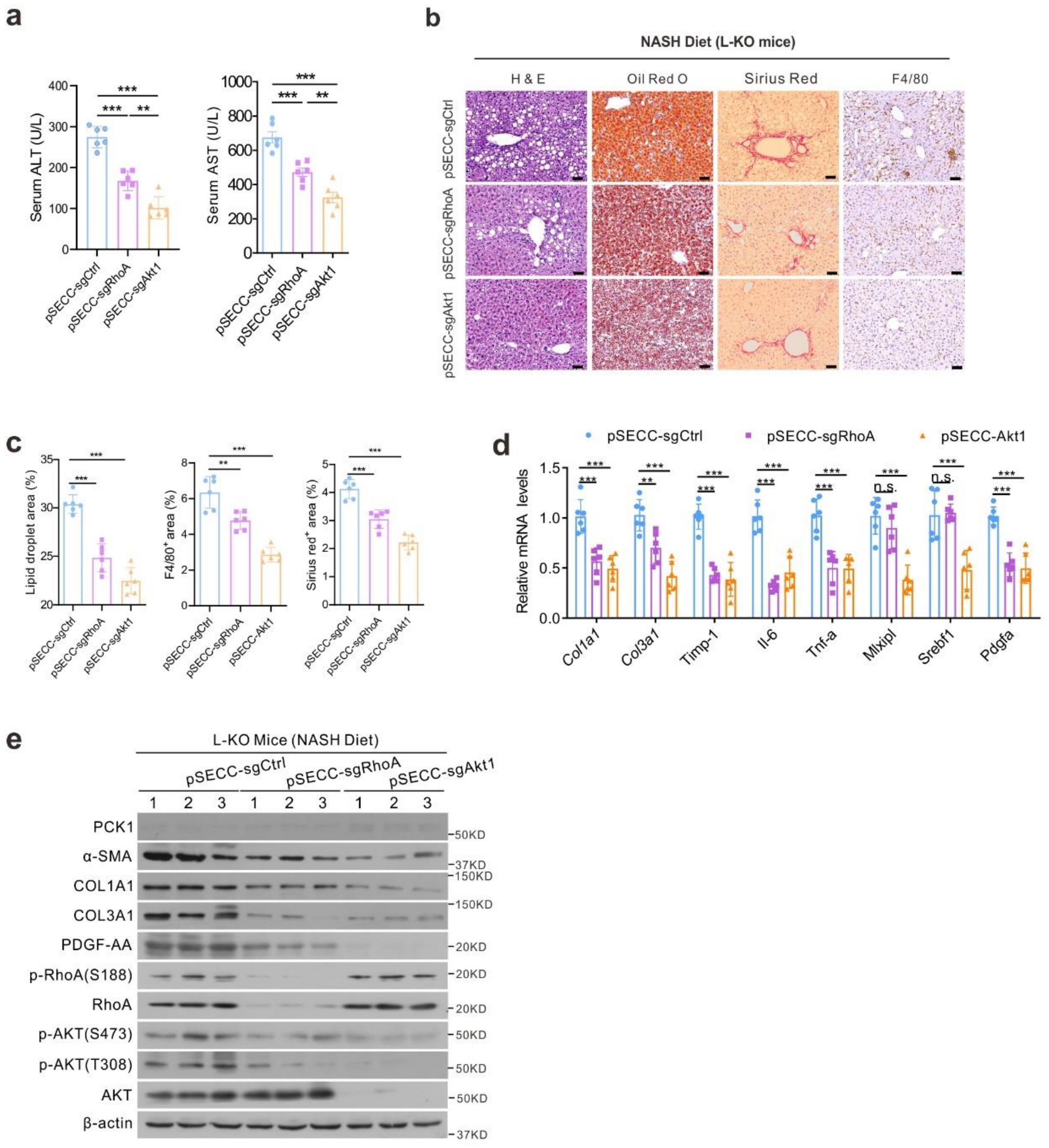


**Supplementary Fig. 7 Genetic inhibition of RhoA or AKT1 protects mice from diet-induced NASH pathologies in L-KO mice. a** Levels of AST and ALT in plasma. **b** Representative HE staining, Oil Red O staining and immunohistochemical staining of liver tissues from L-KO mice infected with the control pSECC-sgTom lentivirus, the pSECC-sgAKT1 or the pSECC-sgRhoA lentivirus. Scale bars: 50 μm. **c** Quantification of the Oil Red O and immunohistochemical staining. **d** Relative mRNA expression in liver tissue of various groups was analyzed by qPCR. **e** Western blot analysis of indicated protein expression in mice liver tissues. Data expressed as mean ± SEM; **P* < 0.05, ** *P* < 0.01, ****P* < 0.001; n.s., not statistically significant. *P* values obtained via one-way ANOVA with Tukey's post hoc test.

### Supplementary Tables

**Supplementary Table 1.** Clinical informations of patients with NASH.

| **Clinical manifestation** | **Patients with NASH** |
|---|---|
| Age, years(mean±SEM) | 44.06 ± 2.74 |
| Gender, *n* (M/F) | 18/18 |
| BMI, kg/$m^2$ | 25.25 ± 1.51 |
| **Laboratory tests**(mean ± SEM) | |
| ALT, U/L | 136.2 ± 11.47 |
| AST, U/L | 74 ± 6.52 |
| ALP, U/L | 105.97 ± 8.50 |
| γ-GT, U/L | 83.17 ± 10.49 |
| TG, mmol/L | 1.41 ± 0.12 |
| TC, mmol/L | 4.56 ± 0.20 |

ALT,Alanine aminotransferase; ALP, Alkaline phosphatase; AST, Aspartate aminotransferase; BMI, Body Mass Index; γ-GT, γ-glutamyl-transferase; TC, Total cholesterol; TG, Triglyceride.

**Supplementary Table 2.** Information on reagents.

| **Name** | **Supplier** | **Cat no.** |
|---|---|---|
| MK2206 | Selleckchem | S1078 |
| Rhosin | MCE | HY-12646 |
| Palmatic acid | Sigma Aldrich | P0500 |
| BSA | Solarbio | A8020-100G |
| pAdTrack-TO4 | Dr.T-C He, University of Chicago | N/A |
| pSEB-3Flag | Dr.T-C He, University of Chicago | N/A |
| AdEasy-BJ5183 E. coli | Lab stock | N/A |
| DH10B Chemically Competent E. coli | Lab stock | N/A |
| AdGFP | Lab stock | N/A |
| AdPCK1 | Lab stock | N/A |
| pSEB-3Flag - PCK1 | Lab stock | N/A |
| Lenticrispr-V2 | Lab stock | N/A |
| Lenticrispr-*PCK1* KO1 | Lab stock | N/A |
| Lenticrispr-*PCK1* KO2 | Lab stock | N/A |

**Supplementary Table 3.** Primer sequences used in this study.

| **Name** | **Sequence** | **Supplier** |
|---|---|---|
| sub-clone primer: *PCK1*-Forward | CGCGGATCCACCATGGGCCCTCCTCAGCTGCAAAACGGCC | TsingKe Biological Technology, China |
| sub-clone primer: *PCK1*-Reverse | CCCAAGCTTCTACATCTGGCTTATTCTTTGCTTCAAG | TsingKe Biological Technology, China |
| real-time PCR: *PCK1*-Forward | ATGGAGGAAGAGGGCATCCT | TsingKe Biological Technology, China |
| real-time PCR: *PCK1*-Reverse | ACGTACATGGTGCGACCTTT | TsingKe Biological Technology, China |
| real-time PCR: *β-ACTIN*-Forward | AGGCCAACCGCGAGAAGATGACC | TsingKe Biological Technology, China |
| real-time PCR: *β-ACTIN*-Reverse | GAAGTCCAGGGCGACGTAGCAC | TsingKe Biological Technology, China |
| real-time PCR: *ACTA2*-Forward | GGGGTGATGGTGGGAATG | TsingKe Biological Technology, China |
| real-time PCR: *ACTA2*-Reverse | GCAGGGTGGGATGCTCTT | TsingKe Biological Technology, China |
| real-time PCR: *COL1A1*- | GACGGCTCAGAGTCACCCA | TsingKe Biological Technology, China |

| | | |
|---|---|---|
| Forward | | |
| real-time PCR: *COL1A1*-Reverse | GGAGACCACGAGGACCAGA | TsingKe Biological Technology, China |
| real-time PCR: *COL3A1*-Forward | GCTCGGGGTAATGACGGT | TsingKe Biological Technology, China |
| real-time PCR: *COL3A1*-Reverse | AGGAATGCCAGCGGGAC | TsingKe Biological Technology, China |
| real-time PCR: *PDGFA*-Forward | TGTCAAGTGCCAGCCCTCC | TsingKe Biological Technology, China |
| real-time PCR: *PDGFA*-Reverse | CCGTGTCCTCTTCCCGATAAT | TsingKe Biological Technology, China |
| real-time PCR: *TIMP1*-Forward | GCTTCTGGCATCCTGTTGTT | TsingKe Biological Technology, China |
| real-time PCR: *TIMP1*-Reverse | TGGTTGACTTCTGGTGTCCC | TsingKe Biological Technology, China |
| real-time PCR: *SERPINH1*-Forward | GCCATGTTCTTCAAGCCACA | TsingKe Biological Technology, China |
| real-time PCR: | CTTTTCCTTCTCGTCGTCGTA | TsingKe Biological |

| | | |
|---|---|---|
| *SERPINH1*-Reverse | | Technology, China |
| real-time PCR: *GFAP*-Forward | GCACGCAGTATGAGGCAATG | TsingKe Biological Technology, China |
| real-time PCR: *GFAP*-Reverse | CCAGGTCGCAGGTCAAGGA | TsingKe Biological Technology, China |
| real-time PCR: *MMP2*-Forward | TTTGACGGTAAGGACGGACTC | TsingKe Biological Technology, China |
| real-time PCR: *MMP2*-Reverse | CCTGGAAGCGGAATGGAAAC | TsingKe Biological Technology, China |
| real-time PCR: *VIM*-Forward | GAGAACTTTGCCGTTGAAGC | TsingKe Biological Technology, China |
| real-time PCR: *VIM*-Reverse | TCCAGCAGCTTCCTGTAGGT | TsingKe Biological Technology, China |
| real-time PCR: *PGC1A*-Forward | GATGGCCTGTTTGATGACAG | TsingKe Biological Technology, China |
| real-time PCR: *PGC1A*-Reverse | TTTGGGTGGTGACACAGAAT | TsingKe Biological Technology, China |
| real-time PCR: *FOXO1*-Forward | TGTCAACCTATGGCAGCCAG | TsingKe Biological Technology, China |
| real-time PCR: *FOXO1*-Reverse | GCAGAGGCACTTGTACAGGT | TsingKe Biological Technology, China |

| | | |
|---|---|---|
| real-time PCR: *ATF3*-Forward | TGAGTGCTTCTGCCATCGTC | TsingKe Biological Technology, China |
| real-time PCR: *ATF3*-Reverse | GGCTACCTCGGCTTTTGTG | TsingKe Biological Technology, China |
| real-time PCR: *HNF4A*-Forward | GCCTACCTCAAAGCCATCAT | TsingKe Biological Technology, China |
| real-time PCR: *HNF4A*-Reverse | CGGTCGTTGATGTAGTCCTC | TsingKe Biological Technology, China |
| real-time PCR: *CEBPB*-Forward | TCGCAGGTCAAGAGCAAGG | TsingKe Biological Technology, China |
| real-time PCR: *CEBPB*-Reverse | GAACAAGTTCCGCAGGGTG | TsingKe Biological Technology, China |
| real-time PCR: *SREBF1*-Forward | TCTGGAGGCATCGCAAGC | TsingKe Biological Technology, China |
| real-time PCR: *SREBF1*-Reverse | CAGCAGGTGACGGATGAGG | TsingKe Biological Technology, China |
| real-time PCR: *NR1H3*-Forward | GGTACAACCCTGGGAGTGAGA | TsingKe Biological Technology, China |
| real-time PCR: *NR1H3*-Reverse | TGGGGATGGTGGATGGAG | TsingKe Biological Technology, China |

| | | |
|---|---|---|
| real-time PCR: *MLXIPL*-Forward | GTCGGCAATGCTGACATGA | TsingKe Biological Technology, China |
| real-time PCR: *MLXIPL*-Reverse | GCTGAAGAGGGAGTCAACCAC | TsingKe Biological Technology, China |
| real-time PCR: *NR1H4*-Forward | AACTCACCCCAGATCAACAGAC | TsingKe Biological Technology, China |
| real-time PCR: *NR1H4*-Reverse | GCTTCAACCGCAGACCCT | TsingKe Biological Technology, China |
| real-time PCR: *Il-6*-Forward | GTTGTGCAATGGCAATTCTGA | TsingKe Biological Technology, China |
| real-time PCR: *Il-6*- Reverse | AAGGACTCTGGCTTTGTCTTTCT | TsingKe Biological Technology, China |
| real-time PCR: *Tnf-α*-Forward | CCTGCCCCAAGGACACC | TsingKe Biological Technology, China |
| real-time PCR: *Tnf-α*-Reverse | AGAGCAATGACTCCAAAGTAGACC | TsingKe Biological Technology, China |
| real-time PCR: *Il-1b*-Forward | AAGCCTCGTGCTGTCGGA | TsingKe Biological Technology, China |
| real-time PCR: *Il-1b* - Reverse | CCATCTTCTTCTTTGGGTATTGC | TsingKe Biological Technology, China |
| real-time PCR: *Il-10*-Forward | GGTTGCCAAGCCTTATCGG | TsingKe Biological Technology, China |

| | | |
|---|---|---|
| real-time PCR: *Il-10*-Reverse | ATTTTCACAGGGGAGAAATCG | TsingKe Biological Technology, China |
| real-time PCR: *Ccl2*-Forward | TGTGCTGACCCCAAGAAGG | TsingKe Biological Technology, China |
| real-time PCR: *Ccl2*- Reverse | GGTGGTTGTGGAAAAGGTAGTG | TsingKe Biological Technology, China |
| real-time PCR: *Acta2*-Forward | CCCTGAAGAGCATCCGACA | TsingKe Biological Technology, China |
| real-time PCR: *Acta2*- Reverse | CATCTCCAGAGTCCAGCACAA | TsingKe Biological Technology, China |
| real-time PCR: *Col1a1*-Forward | ACCCTGCCCGCACATG | TsingKe Biological Technology, China |
| real-time PCR: *Col1a1*- Reverse | CCCTCGCTTCCGTACTCG | TsingKe Biological Technology, China |
| real-time PCR: *Lama2*-Forward | TCCAGCCAAACCATCAGTCC | TsingKe Biological Technology, China |
| real-time PCR: *Lama2*- Reverse | CCACAAGAAGGTCCAATCCAAC | TsingKe Biological Technology, China |
| real-time PCR: *Timp-1*-Forward | CCCAGAAATCAACGAGACCA | TsingKe Biological Technology, China |
| real-time PCR: *Timp-1*-Reverse | ACGCCAGGGAACCAAGAA | TsingKe Biological Technology, China |

| | | |
|---|---|---|
| real-time PCR: *Col3a1*-Forward | GCCCACAGCCTTCTACACCT | TsingKe Biological Technology, China |
| real-time PCR: *Col3a1*-Reverse | TCCCGGATAGCCACCCA | TsingKe Biological Technology, China |
| real-time PCR: *Acly*-Forward | CTCATTGAACCCTTCGTCCC | TsingKe Biological Technology, China |
| real-time PCR: *Acly* - Reverse | CCTCGGTATTCAGCTTTTCGT | TsingKe Biological Technology, China |
| real-time PCR: *Fasn*-Forward | CTGCCTCCGTGGACCTTATC | TsingKe Biological Technology, China |
| real-time PCR: *Fasn* - Reverse | GCACAGACACCTTCCCGTCA | TsingKe Biological Technology, China |
| real-time PCR: *Acaca*–Forward | ATTGCCTATGAACTCAACAGCG | TsingKe Biological Technology, China |
| real-time PCR: *Acaca* - Reverse | TGACAAGGTGGCGTGAAGG | TsingKe Biological Technology, China |
| real-time PCR: *Scd1*-Forward | TTCCTTATCATTGCCAACACCA | TsingKe Biological Technology, China |
| real-time PCR: *Scd1*- Reverse | TCGCCCCAGCAGTACCAG | TsingKe Biological Technology, China |
| real-time PCR: *Elovl6*-Forward | TGCGGGCTGCGGGTTT | TsingKe Biological Technology, China |

| | | |
|---|---|---|
| real-time PCR: *Elovl6*- Reverse | GCCTTCGTGGCTTTCTTCACT | TsingKe Biological Technology, China |
| real-time PCR: *Cpt1a*-Forward | TGTCCAAGTATCTGGCAGTCG | TsingKe Biological Technology, China |
| real-time PCR: *Cpt1a* - Reverse | CATAGCCGTCATCAGCAACC | TsingKe Biological Technology, China |
| real-time PCR: *Mlxipl*-Forward | GCTGCGGGATGAAATAGAGG | TsingKe Biological Technology, China |
| real-time PCR: *Mlxipl*-Reverse | TCAAATAAAGGTCGGATGAGGA | TsingKe Biological Technology, China |
| *real-time PCR:* *Srebf1-Forward* | TTCTGGAGACATCGCAAACAA | TsingKe Biological Technology, China |
| *real-time PCR:* *Srebf1- Reverse* | TGGTAGACAACAGCCGCATC | TsingKe Biological Technology, China |
| real-time PCR: *Ppara*-Forward | GACATTTCCCTGTTTGTGGCT | TsingKe Biological Technology, China |
| real-time PCR: *Ppara*-Reverse | GCTGCGTCGGACTCGGT | TsingKe Biological Technology, China |
| real-time PCR: *Cd36*-Forward | ACTGTGGGCTCATTGCTGG | TsingKe Biological Technology, China |
| real-time PCR: *Cd36*-Reverse | TGATTTTGCTGCTGTTCTTTGC | TsingKe Biological Technology, China |

| | | |
|---|---|---|
| real-time PCR: *Slc27a1*-Forward | GCTCCTGCGGCTTCAACA | TsingKe Biological Technology, China |
| real-time PCR: *Slc27a1*-Reverse | GCGCTATCGCCCTTTCG | TsingKe Biological Technology, China |
| real-time PCR: *Cidea*-Forward | CCTGGTTACGCTGGTGCTG | TsingKe Biological Technology, China |
| real-time PCR: *Cidea*-Reverse | TGGCTATTCCCGATTTCTTTG | TsingKe Biological Technology, China |
| real-time PCR: *Cidec*-Forward | AAGGTTCGCAAAGGCATCA | TsingKe Biological Technology, China |
| real-time PCR: *Cidec*-Reverse | GGCTTCTGGGAAAGGGCTA | TsingKe Biological Technology, China |
| real-time PCR: *Yap1*-Forward | TTTCGGCAGGCAATACGG | TsingKe Biological Technology, China |
| real-time PCR: *Yap1*-Reverse | GGGTGCTTTGGCTGATGGT | TsingKe Biological Technology, China |
| real-time PCR: *Ctgf*-Forward | TTGGCCCAGACCCAACTA | TsingKe Biological Technology, China |
| real-time PCR: *Ctgf* -Reverse | GCAGGAGGCGTTGTCATT | TsingKe Biological Technology, China |
| real-time PCR: *Gli1*-Forward | CGTTTGAAGGCTGTCGGAAGT | TsingKe Biological Technology, China |

| | | |
|---|---|---|
| real-time PCR: *Gli1*-Reverse | GCGGAGCGAGCTGGGAT | TsingKe Biological Technology, China |
| real-time PCR: *Tgfb1*-Forward | CCGCAACAACGCCATCTA | TsingKe Biological Technology, China |
| real-time PCR: *Tgfb1*-Reverse | ACTGCCGTACAACTCCAGTGAC | TsingKe Biological Technology, China |
| real-time PCR: *Pdgfa*-Forward | TGTAACACCAGCAGCGTCAA | TsingKe Biological Technology, China |
| real-time PCR: *Pdgfa*-Reverse | CCTTCCTGTCTCCTCCTCCC | TsingKe Biological Technology, China |
| real-time PCR: *Igf1*-Forward | GGACCGAGGGGCTTTTACT | TsingKe Biological Technology, China |
| real-time PCR: *Igf1*-Reverse | ATAGAGCGGGCTGCTTTTG | TsingKe Biological Technology, China |
| real-time PCR: Vegfa-Forward | CTACTGCCGTCCGATTGAGA | TsingKe Biological Technology, China |
| real-time PCR: *Vegfa*-Reverse | CTGGCTTTGGTGAGGTTTGAT | TsingKe Biological Technology, China |
| real-time PCR: *Fgf21*-Forward | GCATACCCCATCCCTGACTC | TsingKe Biological Technology, China |
| real-time PCR: *Fgf21*-Reverse | GGCTGTTGGCAAAGAAACCTA | TsingKe Biological Technology, China |

| | | |
|---|---|---|
| real-time PCR: *Fgfr1*-Forward | GGATTCTGTGGTGCCTTCTGA | TsingKe Biological Technology, China |
| real-time PCR: *Fgfr1*-Reverse | TTGTCTGGCCCGATCTTACTC | TsingKe Biological Technology, China |
| real-time PCR: *Myc*-Forward | GACTGTATGTGGAGCGGTTTCT | TsingKe Biological Technology, China |
| real-time PCR: *Myc*-Reverse | TCGTTGAGCGGGTAGGGA | TsingKe Biological Technology, China |
| real-time PCR: *Col5a2*-Forward | TGTGCGGGGCAGTGTAGG | TsingKe Biological Technology, China |
| real-time PCR: *Col5a2*-Reverse | TCCCAGGGTCTGTTTTGTTTG | TsingKe Biological Technology, China |
| real-time PCR: *Col6a1*-Forward | GGGGTCAAAGGGGCAAAG | TsingKe Biological Technology, China |
| real-time PCR: *Col6a1*-Reverse | GGCAATCTCAAAGTTCTGTAGGC | TsingKe Biological Technology, China |
| real-time PCR: *Lpar1*-Forward | TCCATACACGAATGAGCAACC | TsingKe Biological Technology, China |
| real-time PCR: *Lpar1*-Reverse | TGGCGAACATAGCCAAAGAT | TsingKe Biological Technology, China |
| real-time PCR: *Lpar3*-Forward | CTGCTCGCACTGCTCAACTC | TsingKe Biological Technology, China |

| | | |
|---|---|---|
| real-time PCR: *Lpar3*-Reverse | CTGGCTGCCCGTCTCG | TsingKe Biological Technology, China |
| real-time PCR: *Gpat3*-Forward | CTTCCAGACAGCAGCCTCAA | TsingKe Biological Technology, China |
| real-time PCR: *Gpat3*-Reverse | TCCCCATCAATCCACCGT | TsingKe Biological Technology, China |
| real-time PCR: *Gpat4*-Forward | CAAGCCCTACACCAACGGAA | TsingKe Biological Technology, China |
| real-time PCR: *Gpat4*-Reverse | TGGCAGGAGGAAGCAATACC | TsingKe Biological Technology, China |
| real-time PCR: *Agpat2*-Forward | CTGCTGTTGCTGCTTGTGC | TsingKe Biological Technology, China |
| real-time PCR: *Agpat2*-Reverse | CCTCCAGTTTCTTCTGTCCG | TsingKe Biological Technology, China |
| real-time PCR: *Agpat3*-Forward | GCTTGCCTACCTGAAGACCC | TsingKe Biological Technology, China |
| real-time PCR: *Agpat3*-Reverse | CCAAACCGCTCGCACATC | TsingKe Biological Technology, China |
| real-time PCR: *Lpin1*-Forward | TGCTCATCCACCAGAGTAAGG | TsingKe Biological Technology, China |
| real-time PCR: *Lpin1*-Reverse | TCCGTGAGGTCGTCCAGAT | TsingKe Biological Technology, China |

| | | |
|---|---|---|
| real-time PCR: *Lpin2*-Forward | CCCCTCCTGGGATTCTGTC | TsingKe Biological Technology, China |
| real-time PCR: *Lpin2*-Reverse | TGAAAAGGCGAGCACTGGTA | TsingKe Biological Technology, China |
| real-time PCR: *Lpin3*-Forward | GGATGACCCAAACCTCGTG | TsingKe Biological Technology, China |
| real-time PCR: *Lpin3*-Reverse | TTGCGGCTTTCTCCCTCT | TsingKe Biological Technology, China |
| real-time PCR: *Dgat1*-Forward | AAGACGGGCGGACCAGC | TsingKe Biological Technology, China |
| real-time PCR: *Dgat1*-Reverse | CACCAGGATGCCATACTTGATA | TsingKe Biological Technology, China |
| real-time PCR: *Dgat2*-Forward | CTGCGGGGTGAGCGTC | TsingKe Biological Technology, China |
| real-time PCR: *Dgat2*-Reverse | ACCTTTCTTGGGCGTGTTC | TsingKe Biological Technology, China |
| real-time PCR: *Mmp1*-Forward | GGCTGAAAGTGACTGGGAAAC | TsingKe Biological Technology, China |
| real-time PCR: *Mmp1*-Reverse | TGGCAAATCTGGCGTGTAA | TsingKe Biological Technology, China |
| real-time PCR: *Mmp9*-Forward | GCCCTGAACCTGAGCCA | TsingKe Biological Technology, China |

| | | |
|---|---|---|
| real-time PCR: *Mmp9*-Reverse | ACTTCCCATCCTTGAACAAATAC | TsingKe Biological Technology, China |
| real-time PCR: *Lama2*-Forward | TCCAGCCAAACCATCAGTCC | TsingKe Biological Technology, China |
| real-time PCR: *Lama2*-Reverse | CCACAAGAAGGTCCAATCCAAC | TsingKe Biological Technology, China |
| real-time PCR: *Lama3*-Forward | CCGCTCGGGCTCCTATT | TsingKe Biological Technology, China |
| real-time PCR: *Lama3*-Reverse | ACATGCTGCTTGCACTGACA | TsingKe Biological Technology, China |
| real-time PCR: *Lama5*-Forward | GGAATATGTCGGTGAGGATTCA | TsingKe Biological Technology, China |
| real-time PCR: *Lama5*-Reverse | TCCAGGTAGAAGATGGCTAGATG | TsingKe Biological Technology, China |
| real-time PCR: *β-Actin*-Forward | CGTTCAATACCCCAGCCATG | TsingKe Biological Technology, China |
| real-time PCR: *β-Actin*-Reverse | GACCCCGTCACCAGAGTCC | TsingKe Biological Technology, China |
| ChIP qPCR: *PCK1*-Forward 1 | CCCAAAGCATAACTGACCCTG | TsingKe Biological Technology, China |
| ChIP qPCR: *PCK1*-Reverse 1 | TTAAATACTGTGGAAAAGAATAGCC | TsingKe Biological Technology, China |

| | | |
|---|---|---|
| ChIP qPCR: *PCK1*-Forward 2 | TGGTTGAGGGCTCGAAGTC | TsingKe Biological Technology, China |
| ChIP qPCR: *PCK1*-Reverse 2 | ACGGCCAGGGTCAGTTATG | TsingKe Biological Technology, China |
| ChIP qPCR: *PCK1*-Forward 3 | CCCAGCATTCATTAACAACTATCT | TsingKe Biological Technology, China |
| ChIP qPCR: *PCK1*-Reverse 3 | TGCTTGGTGGCAGAACCTC | TsingKe Biological Technology, China |
| sgRNA: *PCK1*-Forward | CACCGGCTGAAGAAGTATGACAAC | TsingKe Biological Technology, China |
| sgRNA: *PCK1*-Reverse | AAACGTTGTCATACTTCTTCAGCC | TsingKe Biological Technology, China |
| TA cloning: *PCK1*-Forward | AACCTGTGGATCTCCCTTC | TsingKe Biological Technology, China |
| TA cloning: *PCK1*-Reverse | CAAATCAATGTTCCGCTCA | TsingKe Biological Technology, China |
| shRNA: *ATF3* -Forward 1 | TGCAAAGTGCCGAAACAAGATTCAAGAG ATCTTGTTTCGGCACTTTGCTTTTTTC | TsingKe Biological Technology, China |
| shRNA: *ATF3* -Reverse 1 | TCGAGAAAAAAGCAAAGTGCCGAAACAA GATCTCTTGAATCTTGTTTCGGCACTTTG CA | TsingKe Biological Technology, China |
| shRNA: | TGAGAAACCTCTTTATCCAATTCAAGAGA | TsingKe Biological |

| | | |
|---|---|---|
| *ATF3* -Forward 2 | TTGGATAAAGAGGTTTCTCTTTTTTC | Technology, China |
| shRNA: *ATF3* -Reverse 2 | TCGAGAAAAAAGAGAAACCTCTTTATCCAATCTCTTGAATTGGATAAAGAGGTTTCTCA | TsingKe Biological Technology, China |
| shRNA: *ATF3* -Forward 3 | TGGACTCCAGAAGATGAGAGTTCAAGAGACTCTCATCTTCTGGAGTCCTTTTTTC | TsingKe Biological Technology, China |
| shRNA: *ATF3* -Reverse 3 | TCGAGAAAAAAGGACTCCAGAAGATGAGAGTCTCTTGAACTCTCATCTTCTGGAGTCCA | TsingKe Biological Technology, China |

**Supplementary Table 4.** The antibody information.

| **Name** | **Supplier** | **Cat no.** | **Clone no.** |
|---|---|---|---|
| PCK1 | Bioworld Technology, USA | BS6870 | Polyclonal |
| PPARα | Proteintech, USA | 15540-1-AP | Polyclonal |
| CD36 | Abcam, USA | Ab133625 | EPR6573 |
| FATP1 | Affinity Biosciences, USA | DF7716 | Polyclonal |
| α-SMA | Cell Signaling Technology, USA | 19245T | D4K9N |
| PDGF-AA | Abcam, USA | Ab216619 | Polyclonal |
| CIDEC | Novus Biologicals, USA | NB100-430SS | Polyclonal |
| β-ACTIN | ZSGB-BIO, China | TA-09 | OTI1 |
| CIDEA | Proteintech, USA | 13170-1-AP | Polyclonal |
| COL1A1 | Abcam, USA | Ab34710 | Polyclonal |
| p-AKT (S473) | Bioworld Technology, USA | BS4007 | Polyclonal |
| p-AKT (T308) | Bioworld Technology, USA | AP0056 | Polyclonal |
| AKT | Bioworld Technology, USA | AP0059 | Polyclonal |
| p-RhoA | Abcam, USA | Ab41435 | Polyclonal |

| | | | |
|---|---|---|---|
| (S188) | | | |
| RhoA | Abcam, USA | Ab187027 | EPR18134 |
| F4/80 | Cell Signaling Technology, USA | 70076T | D2S9R |
| ATF3 | Abcam, USA | Ab207434 | EPR19488 |
| COL3A1 | Proteintech | 22734-1-AP | Polyclonal |
| RhoA | Cell Signaling Technology, USA | 2117T | 67B9 |
| RhoB | Cell Signaling Technology, USA | 2098T | Polyclonal |
| RhoC | Cell Signaling Technology, USA | 3430T | D40E4 |
| RAC1/2/3 | Cell Signaling Technology, USA | 2465T | Polyclonal |
| p-RAC1 | Cell Signaling Technology, USA | 2461T | Polyclonal |
| CDC42 | Cell Signaling Technology, USA | 2466T | 11A11 |
| PI3 Kinase p85 | Cell Signaling Technology, USA | 4257 | 19H8 |
